# DNA sequence is a major determinant of tetrasome dynamics

**DOI:** 10.1101/629485

**Authors:** O. Ordu, A. Lusser, N. H. Dekker

## Abstract

Eukaryotic genomes are hierarchically organized into protein-DNA assemblies for compaction into the nucleus. Nucleosomes, with the (H3-H4)_2_ tetrasome as a likely intermediate, are highly dynamic in nature by way of several different mechanisms. We have recently shown that tetrasomes spontaneously change the direction of their DNA wrapping between left- and right-handed conformations, which may prevent torque build-up in chromatin during active transcription or replication. DNA sequence has been shown to strongly affect nucleosome positioning throughout chromatin. It is not known, however, whether DNA sequence also impacts the dynamic properties of tetrasomes. To address this question, we examined tetrasomes assembled on a high-affinity DNA sequence using freely orbiting magnetic tweezers. In this context, we also studied the effects of mono- and divalent salts on the flipping dynamics. We found that neither DNA sequence nor altered buffer conditions affect overall tetrasome structure. In contrast, tetrasomes bound to high-affinity DNA sequences showed significantly altered flipping kinetics, predominantly via a reduction in the lifetime of the canonical state of left-handed wrapping. Increased mono- and divalent salt concentrations counteracted this behaviour. Thus, our study indicates that high-affinity DNA sequences impact not only the positioning of the nucleosome, but that they also endow the subnucleosomal tetrasome with enhanced conformational plasticity. This may provide a means to prevent histone loss upon exposure to torsional stress, thereby contributing to the integrity of chromatin at high-affinity sites.

**STATEMENT OF SIGNIFICANCE:** Canonical (H3-H4)_2_ tetrasomes possess high conformational flexibility, as evidenced by their spontaneous flipping between states of left- and right-handed DNA wrapping. Here, we show that these conformational dynamics of tetrasomes cannot be described by a fixed set of rates over all conditions. Instead, an accurate description of their behavior must take into account details of their loading, in particular the underlying DNA sequence. *In vivo*, differences in tetrasome flexibility could be regulated by modifications of the histone core or the tetrasomal DNA, and as such constitute an intriguing, potentially adjustable mechanism for chromatin to accommodate the torsional stress generated by processes such as transcription and replication.

## INTRODUCTION

The nucleosome is the basic complex of chromatin in eukaryotic cells (1-3). It comprises 147 base pairs (bp) of deoxyribonucleic acid (DNA) wrapped around a disk-shaped assembly of eight histone proteins in ∼1.7 turns (4-6). The histone octamer consists of two copies of two types of heterodimers, one of which is formed by histones H3 and H4, and the other by histones H2A and H2B (7,8). During nucleosome formation, first the two H3-H4 dimers assemble onto the DNA to form a tetrasome, after which the binding of two H2A/H2B dimers completes the full nucleosome (9). The resulting compaction upon DNA wrapping limits the accessibility of the genome for essential cellular processes, such as transcription, replication and repair. Hence, the positioning and stability of nucleosomes play a key role in gene regulation and cellular function.

Recent research has provided significant new insights into the structure and especially the dynamics of nucleosomes (10). For example, studies using single-molecule techniques have revealed the intrinsically dynamic nature of nucleosomes in the form of ‘breathing’, i.e. the transient un- and rewrapping of the nucleosomal DNA ends (11-14). Nucleosomes were also found to ‘gap’ by transiently opening and closing the two turns of nucleosomal DNA along the direction of the superhelical axis (15). The structure, stability and dynamics of nucleosomes have furthermore been shown to be extensively regulated by post-translational modifications (16), chaperones (17), and ATP-dependent remodelers (18). Furthermore, nucleosome composition and dynamics can be altered by forces and torques generated by molecular motors that process the genome (19,20). Such external influences, as well as changes in the ambient conditions, can cause nucleosomes to reorganize into different (sub)structures (21). Specifically, tetrasomes, which wrap ∼80 bp of DNA around the (H3-H4)_2_ tetramer (**Figure 1A**), have been observed as stable intermediates in several studies (22-28). Similar to nucleosomes, tetrasomes predominantly adopt a left-handed DNA wrapping, but interestingly, right-handed DNA wrapping has also been identified (29-34). Recently, we showed that tetrasomes can spontaneously switch between these two conformations without unbinding from the DNA (35-37) (**Figure 2A**). This conformational flexibility could mitigate the build-up of torsional stress in chromatin during e.g. transcription or replication and thereby prevent the eviction of histones from the DNA.

**Figure 1.**
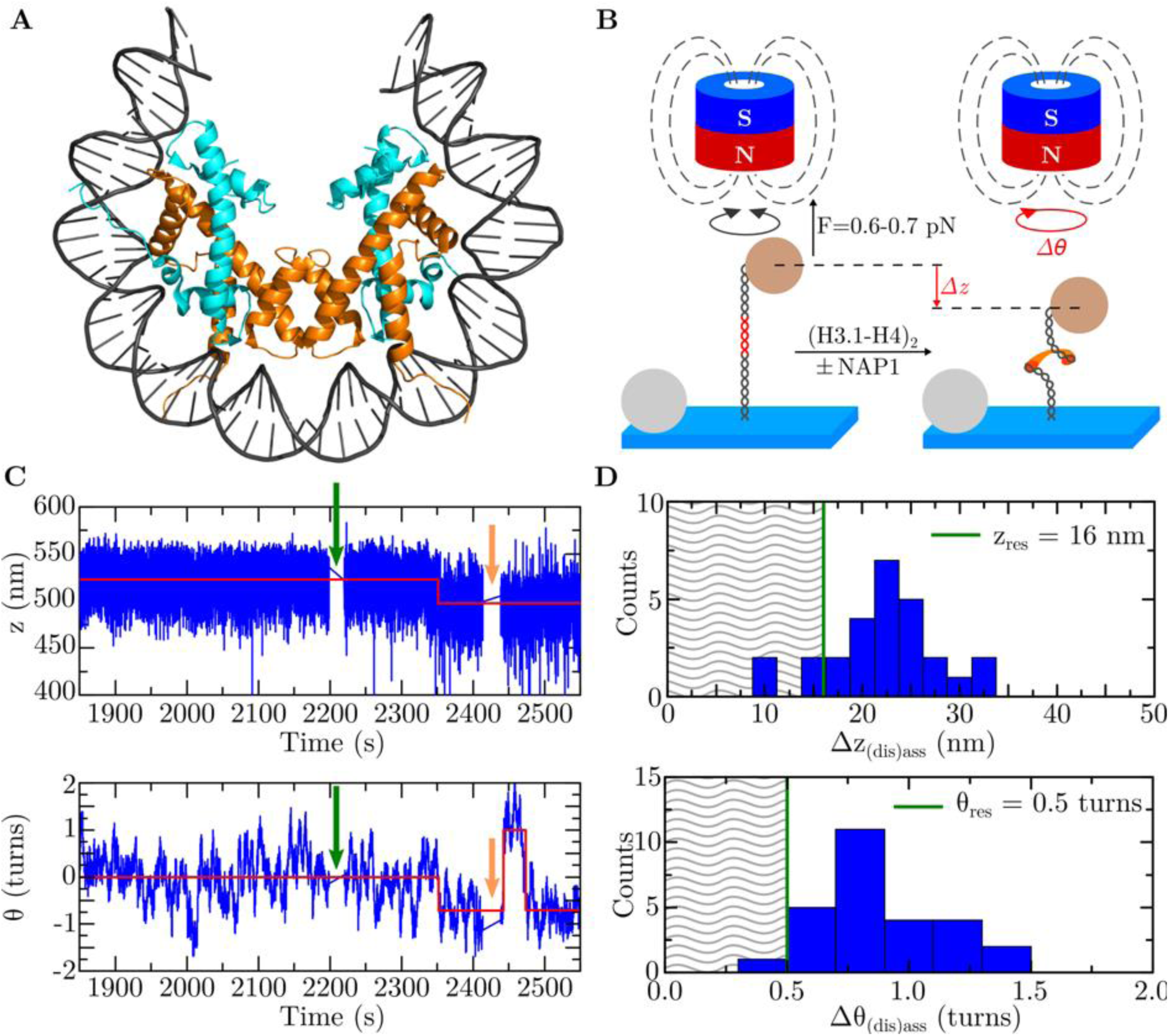
Real-time assembly of single tetrasomes onto DNA with a 601-sequence. **A** Top view on about 80 bp of DNA (dark gray) bound to a tetrameric protein core consisting of the histones H3 (orange) and H4 (cyan). This image was created by modifying the structural data of the *Drosophila* nucleosome from the RCSB Protein Data Bank (PDB) with the PDB identification code 2PYO (82). **B** Experimental assay based on FOMT (49). A single DNA construct (black) containing a 601-sequence (red) at its center (DNA_w/601_) is attached to a coverslip (light blue) at one end, and tethered to a superparamagnetic bead (light brown) at the other end. Above the flow cell, a cylindrically shaped permanent magnet (dark blue/red), with its axis precisely aligned with the DNA tether, exerts a constant force (*F*) on the bead while allowing its free rotation in the (*x,y*)-plane (indicated by the black circular arrow). This enables the direct measurement of the DNA molecule’s length *z* and linking number *Θ*, which upon the assembly of a tetrasome (orange) are changed by *Δz* and *ΔΘ* (indicated by the red straight and circular arrows, respectively). Tetrasome assembly is induced by flushing in either (H3.1-H4)_2_ tetramers alone or pre-incubated histone/NAP1 complexes. Non-magnetic beads attached to the flow cell surface serve as reference for drift correction. This figure is adapted from Ref. (37) (https://doi.org/10.1063/1.5009100). **C** Partial time traces of the length *z* (in nm, top panel) and the linking number *Θ* (in turns, bottom panel) of a DNA_w/601_ molecule before and upon the assembly of a (H3.1-H4)_2_ tetrasome. The formation of a tetrasome simultaneously decreased both quantities in the form of a step identified using a step-finder algorithm (red lines) (**Materials and Methods**). About 60 s after assembly, free proteins were flushed out with measurement buffer (orange arrows) to prevent further histone binding. In this particular experiment, a tetrasome was assembled by flushing in NAP1/histone complexes (green arrows) in buffer A (**Table I**). A corresponding time trace of a DNA_w/601_ molecule upon the assembly of a tetrasome from histone tetramers only is shown in **Supplementary Figure S4. D** Histogram of the changes in DNA length upon assembly and disassembly *Δz*_*(dis)ass*_ (in nm, top panel) of single (H3.1-H4)_2_ tetrasomes (*N*=27) and in DNA linking number upon assembly and disassembly *ΔΘ*_*(dis)ass*_ (in turns, bottom panel) of single (H3.1-H4)_2_ tetrasomes (*N*=27) in all buffers (**Table I**), shown together with the mean spatial resolution corresponding to the average 1 STD (16 nm and 0.5 turns; green lines) from all experiments. Data beyond the resolution limits (hatched area) was excluded from the determination of the mean change *Δz*_*(dis)ass*_ = 23 ± 5 nm (*n*=25) and *ΔΘ*_*(dis)ass*_ = 0.9 ± 0.2 turns (*n*=26).

**Figure 2.**
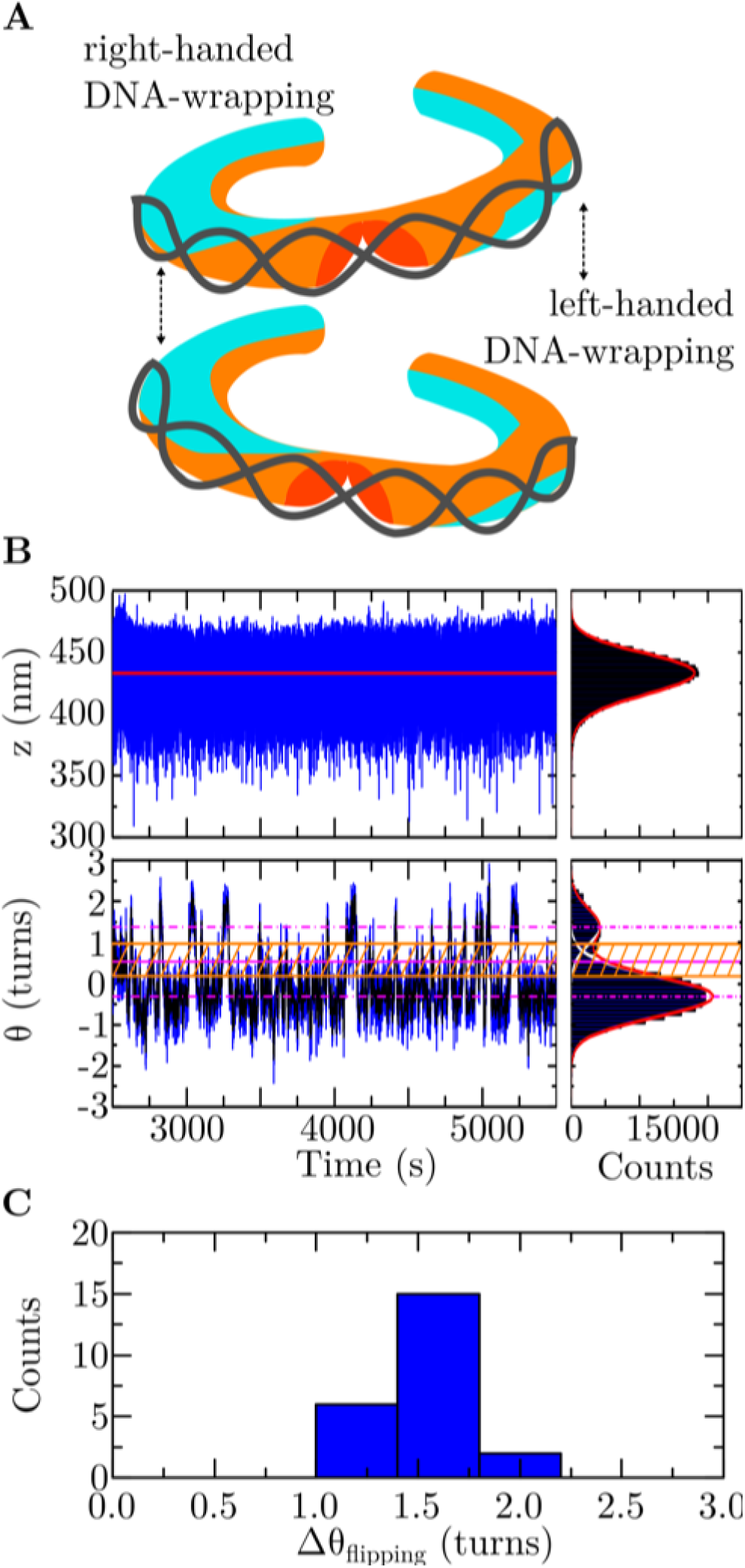
The structural dynamics of single tetrasomes on DNA with a 601-sequence. **A** Representation of the two tetrasome conformations with the DNA wrapped in either a left-handed or a right-handed superhelix. Tetrasomes were observed to spontaneously flip between these two states (35-37). This image is adapted from Ref. (37) (https://doi.org/10.1063/1.5009100). **B** Partial time traces of the length *z* (in nm, top panel) and the linking number *Θ* (in turns, bottom panel) of a DNA molecule after the assembly of a (H3.1-H4)_2_ tetrasome in buffer A (**Table I**) upon flushing in histone/NAP1 complexes. As indicated from the fit by the step-finder algorithm to the time trace (red line, left panel) and the fit of a mirrored gamma function (red line in histogram plot, right panel) to the skewed data, the DNA length remains constant. The DNA linking number spontaneously fluctuates, i.e. ‘flips’, between two states identified by fitting two Gaussian functions (white lines in histogram plot, right panel) underlying the full profile (red line in histogram plot, right panel). The two states correspond to a prevalent left-handed and a less adopted right-handed conformation of DNA wrapping, with the respective mean linking numbers *Θ*_*left*_ = −0.31 ± 0.01 turns and *Θ*_*right*_ = +1.38 ± 0.06 turns (dashed-dotted magenta lines, 95% confidence level for estimated values). Due to drift, the mean value for *Θ*_*left*_ obtained here is offset from the average change in DNA linking number upon tetrasome dis-/assembly (**Figure 1D**, bottom panel). The structural dynamics were quantified in terms of the dwell times in the two states based on a threshold zone (hatched orange area) that is bounded by 1 STD from each mean value (orange solid lines) about their average (solid magenta line) (**Materials and Methods**). A corresponding partial time trace of a DNA_w/601_ molecule upon the assembly of a tetrasome from histone tetramers only is shown in **Supplementary Figure S7**. **C** Histogram of the change in DNA linking number *ΔΘ*_*flipping*_ (in turns) upon flipping of single (H3.1-H4)_2_ tetrasomes in their handedness of DNA-wrapping in all buffers (**Table I**). The data yields a mean value of *ΔΘ*_*flipping*_ = 1.6 ± 0.2 turns (*N*=23). The individual values are provided in **Supplementary Table SIX**.

**Table I.** Composition of the measurement buffers used in this work.

| Measurement buffers | Buffer Components |
| --- | --- |
| **A [Standard]** | **25 mM 4-(2-hydroxyethyl)-1-piperazineethane-sulfonic acid-KOH (HEPES-KOH (pH 7.5); Sigma-Aldrich, St. Louis, MO, United States), 0.1 mM ethylenediaminetetraacetic acid (EDTA; Sigma-Aldrich), 50 mM KCl (Merck, Darmstadt, Germany), 0.01% (w/v) BSA (Sigma-Aldrich), 0.025% (w/v) PEG (Sigma-Aldrich), 0.025% (w/v) PVA (Sigma-Aldrich)** |
| **B** | **10 mM HEPES (pH 7.5), 100 mM KCl, 0.2% (w/v) BSA, 0.1% (w/v) Tween-20 (Sigma-Aldrich), 10 mM NaN<sub>3</sub> (Merck)** |
| **C** | **10 mM HEPES (pH 7.5), 100 mM KCl, 0.2% (w/v) BSA, 0.1% (w/v) Tween-20, 10 mM NaN<sub>3</sub>, 2 mM MgCl<sub>2</sub>** |

DNA sequence has been suggested to play an important role in determining the structural and dynamic properties of nucleosomes (38). The formation of a nucleosome is significantly affected by the ability of the underlying DNA to bend sharply (39,40). This feature has not only been examined in numerous *in vitro* studies (41), but it has also been observed at a global level *in vivo* (42). For example, in yeast it was shown that ∼50% of the *in vivo* nucleosome positions can be attributed to sequence-specific features. In particular, high-affinity nucleosome binding sequences were found to be enriched in intergenic regions where an integral chromatin structure is important, while low affinity sequences were most dominant in highly transcribed regions, such as rRNA and tRNA genes (42). High-affinity sequences were also detected in strongly regulated genes, and it was proposed that such sequences may favour rapid reassembly of nucleosomes promoting repression of these genes under unfavourable conditions (42).

While structure and dynamics of high-affinity nucleosomes have also been studied extensively by *in vitro* single-molecule assays (43-48), an interesting open question is whether association of histones with high-affinity DNA sequences affects the dynamics of subnucleosomal structures, such as (H3-H4)_2_ tetrasomes, which can arise in the course of transcription or other torque-generating processes. In this work we have addressed this question by a single molecule approach. Using freely-orbiting magnetic tweezers (FOMT) (49), we systematically investigated the structure and handedness dynamics of single (H3.1-H4)_2_ tetrasomes assembled onto the well-characterized high-affinity Widom 601 nucleosome-binding sequence (50). In addition, we have examined the contribution of different buffers commonly employed in chromatin studies (51-53) to the dynamics of high-affinity tetrasomes. Our results revealed intriguing effects of DNA sequence on handedness flipping dynamics that can be counteracted by mono- and divalent salts. Thus, our findings provide conceptual evidence for a model in which chromatin regions containing high-affinity nucleosomes may be better protected than other regions from complete histone loss during torque-generating processes, such as transcription or replication, due to inherently increased handedness flipping dynamics of (H3-H4)_2_ tetrasomes.

## MATERIALS AND METHODS

### Preparation of DNA molecules

Tetrasome assembly was performed on linear double-stranded DNA (dsDNA) fragments of 1.96 kilo-base pair (kbp) length containing a single 601-sequence of 147 bp at their center. The two ends of the DNA fragments were ligated to digoxigenin-coated (Roche Diagnostics, Basel, Switzerland) or biotin-coated (Roche Diagnostics) double-stranded fragments (handle) of 643 bp length, respectively, for immobilization and tethering. The schematics, sequence, and preparation of the DNA molecules are described in **Supplementary Figure S1** and **Supplementary Table SI**. Further details on the handles have been described in Ref. (37).

### Expression and purification of proteins

The recombinant canonical *Drosophila* histones H3.1-H4 and nucleosome assembly protein-1 (NAP1) chaperones were expressed and purified as described in Ref. (35) and Ref. (37). While *in vivo* NAP1 has been identified as a chaperone for histones H2A and H2B (54), *in vitro* it has been found to also interact with histones H3 and H4 and prevent histone aggregation (35-37,55-60).

### Preparation of histones and tetrasome assembly

Individual tetrasomes were assembled and monitored in real time using magnetic tweezers (see below) in three buffers with varying compositions of core components employed in previous studies (35-37,51-53) (**Table I**). To allow for direct comparison to our previous study (35), the protein samples were prepared similarly, by incubating 51 nanomolar (nM) of an equimolar solution of H3.1-H4 histones – either without or with 192 nM NAP1 – on ice for 30 min. The incubation buffers were identical to the measurement buffers (**Table I**), except for buffer A. In that case, the incubation buffer contained 0.25% (w/v) Polyethylene Glycol (PEG) and 0.25% (w/v) Polyvinyl Alcohol (PVA), and 0.1% (w/v) Bovine Serum Albumin (BSA), as in our previous study (35). The incubated protein solutions were then diluted by at most 1:5000, and 100 μl of the diluted solution was flushed into the flow cell to assemble a single tetrasome. About 60 s after assembly, 100-300 μl of measurement buffer was flushed through the flow cell to remove free proteins and to prevent further assembly of tetrasomes.

The details of the sample and flow cell preparation for magnetic tweezers experiments have been described in Ref. (37). Where the concentrations and volumes of DNA and magnetic bead solutions used in this work differ, this is stated in **Supplementary Table SII**. Five (out of total *N*_*exp,total*_=20) experiments were performed by resuming the measurement on a tetrasome that did not disassemble during a preceding experiment. Two such experiments involved an exchange between buffers B and C (**Table I**) with 300-500 μl of the respective measurement buffer: In one of these two experiments, a measurement was resumed in buffer C on a tetrasome that had been assembled before in buffer B, and in the other the reverse procedure was employed.

### Magnetic tweezers instrumentation

The assembly and dynamics of an individual tetrasome was monitored for up to 10 h by directly measuring the length and angular position, i.e. the linking number, of a single DNA molecule in each experiment using FOMT (49) (**Figure 1B**). The magnetic tweezers hardware is detailed in Ref. (37). All experiments were performed at an exerted force of 0.6-0.7 piconewton (pN) and at room temperature (22 ºC).

### Data analysis

The analysis of the acquired data was performed using custom-written scripts with built-in functions in Matlab (Mathworks, Natick, MA, USA). An improved version of the custom-written step-finder algorithm described in Ref. (61) was used to detect stepwise changes in the time traces of the DNA length and DNA linking number. Subsequently, these fits were analyzed to identify simultaneous steps in both DNA length and DNA linking number within a time window of 19.1 s (see below), resulting from the assembly or disassembly of a tetrasome (**Figure 1C, Supplementary Figure S4**). With this approach, 48% (*n*=13) of all such steps (*N*=27) were automatically identified, while the remaining 52% (*n*=14) were corrected manually to better match the data. The retrieved sizes of these simultaneous changes in DNA length (*N*=27) and DNA linking number (*N*=27) form the basic quantities that characterize tetrasomes structure (**Supplementary Tables SIII** and **SIV)**. Step sizes were also compared to the mean spatial resolution, determined by averaging the standard deviations (STDs) of the time trace profiles in all experiments (**Figure 1D, Supplementary Figure S5**).

The dynamics of tetrasome structure in terms of handedness flipping (**Figure 2A**) was separately analyzed in the parts of the time traces with a stable baseline in DNA length and DNA linking number (**Figure 2B, Supplementary Figure S7**). The change in DNA linking number upon flipping was determined by fitting a double Gaussian function to the DNA linking number data (*N*=23; **Figure 2B, Supplementary Table SIX**, and **Supplementary Figure S7**) and calculating the difference between the mean values. The probability for a tetrasome to occupy one of the two states was calculated from the relative ratio of the corresponding peak areas of the two Gaussian fits (**Supplementary Table SXI**).

The handedness dynamics of single, flipping (H3.1-H4)_2_ tetrasomes was further analyzed in terms of the times spent in the left- or right-handed state using another custom-written algorithm based on Ref. (62). A threshold zone was defined by 1 STDs from the mean values of the two Gaussian distributions about their average (**Figure 2B, Supplementary Figure S7**). The details of the dwell-time analysis have been described in Ref. (37). Here, the corresponding DNA linking number data of the time traces of sufficiently long duration (>1600 s) were smoothed by filtering over *N*=330 (*N*=1910) points, corresponding to a time average of *τ*_*short*_ = 3.3 s (*τ*_*long*_ = 19.1 s). These time scales were obtained from autocorrelation analysis of the bead’s angular fluctuations, as described in Refs. (37,49) (**Supplementary Figure S2**). The mean dwell time in each state was then determined by combining all datasets obtained in identical buffer conditions and fitting these to an exponential function (**Table II, Figure 3A-C, Supplementary Table SXII**, and **Supplementary Figure S8A-C**). For direct comparison, we also performed dwell-time analysis with the same settings on the partial time traces (*N*=6) of one of our earlier experiments published in Ref. (35).

**Table II.** Summary of quantified properties for NAP1-loaded tetrasomes. Specific buffers employed are indicated within brackets. The results for spontaneously loaded tetrasomes are provided in **Supplementary Table SXII**.

| Quantity | (H3.1-H4)<sub>2</sub> tetrasomes on<br>DNA<sub>random</sub> | (H3.1-H4)<sub>2</sub> tetrasomes on<br>DNA<sub>w/601</sub> |
| --- | --- | --- |
| $\Delta z_{(dis)ass}$ (nm) | 24 ($\pm 3$)<sup>*</sup> | 23 ($\pm 5$)<sup>x</sup> |
| $\Delta \theta_{(dis)ass}$ (turns) | 0.73 ($\pm 0.05$)<sup>*</sup> | 0.9 ($\pm 0.2$)<sup>x</sup> |
| $\Delta \theta_{flipping}$ (turns) | 1.7 ($\pm 0.1$)<sup>*</sup> | 1.6 ($\pm 0.2$)<sup>x</sup> |
| $\tau_{D,left}$ (s) | 183 (+15/-12)<sup>+</sup> [A] | 37 ($\pm 2$)<sup>**</sup> [A] |
| with time average in filtering of<br>3.3 s [N=330] | | 62 ($\pm 3$)<sup>**</sup> [B] |
|  |  | 141 (+11/-10)<sup>**</sup> [C] |
| $\tau_{D,left}$ (s) | 417 (+60/-48)<sup>+</sup> [A] | 67 (+5/-4)<sup>**</sup> [A] |
| with time average in filtering of<br>19.1 s [N=1910] |  | 121 (+8/-7)<sup>**</sup> [B] |
|  |  | 231 (+24/-20)<sup>**</sup> [C] |
| $\tau_{D,right}$ (s) | 14 ($\pm 1$) [A] | 28 (+2/-1)<sup>**</sup> [A] |
| with time average in filtering of<br>3.3 s [N=330] | | 33 ($\pm 1$)<sup>**</sup> [B] |
| | | 30 ($\pm 2$)<sup>**</sup> [C] |
| $\tau_{D,right}$ (s) | 34 (+5/-4)<sup>**</sup> [A] | 48 (+4/-3)<sup>**</sup> [A] |
| with time average in filtering of<br>19.1 s [N=1910] | | 64 ($\pm 4$)<sup>**</sup> [B] |
|  |  | 44 (+5/-4)<sup>**</sup> [C] |
| $\tau_{D,left}/\tau_{D,right}$ | 13 ($\pm 1$)<sup>+</sup> [A] | 1 ($\pm 0$)<sup>+</sup> [A] |
| with time average in filtering of<br>3.3 s [N=330] | | 2 ($\pm 0$)<sup>+</sup> [B] |
| | | 4 ($\pm 0$)<sup>+</sup> [C] |
| $\tau_{D,left}/\tau_{D,right}$ | 12 ($\pm 3$)<sup>+</sup> [A] | 1 ($\pm 0$)<sup>+</sup> [A] |
| with time average in filtering of<br>19.1 s [N=1910] | | 2 ($\pm 0$)<sup>+</sup> [B] |
| | | 5 ($\pm 1$)<sup>+</sup> [C] |
| $\Delta E_{dwell-times}$ (k<sub>B</sub>T) | 2.6 ($\pm 0.1$)<sup>+</sup> [A] | 0.3 ($\pm 0.1$)<sup>+</sup> [A] |
| | | 0.6 ($\pm 0.1$)<sup>+</sup> [B] |
| | | 1.5 ($\pm 0.1$)<sup>+</sup> [C] |
| $\Delta E_{peak-areas}$ (k<sub>B</sub>T) | - | 0.9 ($\pm 0.7$)<sup>+</sup> [A] |
| | | 1.4 ($\pm 0.8$)<sup>+</sup> [B] |
| | | 1.7 (± ----) [C] |

\*Values taken from Ref. (35)
<sup>x</sup>Values obtained for spontaneously loaded tetrasomes on DNA<sub>w/601</sub> are included for improved statistics
<sup>+</sup>Errors calculated by error propagation
<sup>\*\*</sup>Errors correspond to 68% confidence interval for estimated values from exponential fits

**Figure 3.**
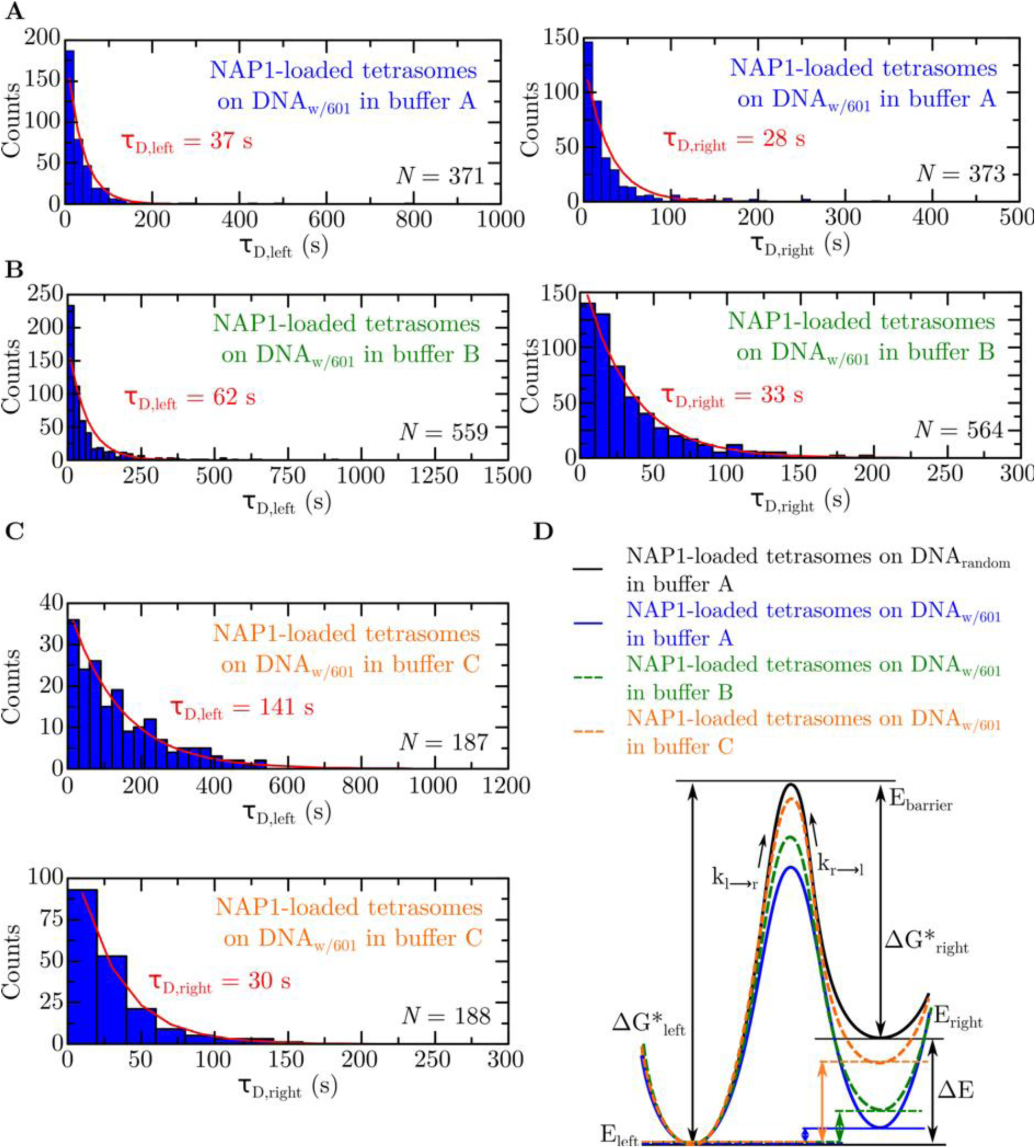
Kinetics and energetics of single tetrasomes on DNA with a 601-sequence. **A** Histograms of the dwell times of a single NAP1-loaded (H3.1-H4)_2_ tetrasome on a DNA_w/601_ molecule in the left-(left panel) and right-handed conformation (right panel) in buffer A (**Tables I** and **II**). Exponential fits (red lines) yielded a mean dwell time of *τ*_*D,left*_ = 37 ± 2 s (*N*=371) and *τ*_*D,right*_ = 28 +2/-1 s (*N*=373), respectively (68% confidence level for estimated values). **B** As in (A), but now in buffer B (**Tables I** and **II**). Exponential fits (red lines) yielded a mean dwell time of *τ*_*D,left*_ = 62 ± 3 s (*N*=559) and *τ*_*D,right*_ = 33 ± 1 s (*N*=564), respectively (68% confidence level for estimated values). **C** Histograms of the dwell times of a single NAP1-loaded (H3.1-H4)_2_ tetrasome on a DNA_w/601_ molecule in the left-handed (top panel) and right-handed conformation (bottom panel) in buffer C (**Tables I** and **II**). Exponential fits (red lines) yielded a mean dwell time of *τ*_*D,left*_ = 141 +11/-10 s (*N*=187) and *τ*_*D,right*_ = 30 ± 2 s (*N*=188), respectively (68% confidence level for estimated values). All data in panels (A)-(C) were obtained by dwell-time analysis of the DNA linking number time traces filtered by averaging over 3.3 s (N=330) (**Materials and Methods**). **D** Schematic energy diagrams of single NAP1-loaded tetrasomes on DNA_w/601_ in all buffer conditions, based on the dwell-time values and the probabilities obtained from the linking number distributions (**Table II, Supplementary Tables SXI, SXIII**-**SXV**). The solid black lines illustrate the energy levels in buffer A (**Table I**) for a NAP1-loaded (H3.1-H4)_2_ tetrasome on DNA_random_, while the solid blue lines depict the energetics for a NAP1-loaded (H3.1-H4)_2_ tetrasome on DNA_w/601_. The green and orange dashed lines show the energy levels for a NAP1-loaded (H3.1-H4)_2_ tetrasome on DNA_w/601_ in buffers B and C, respectively. The free energy differences (*ΔE)* between the left- and right-handed conformations of tetrasomes on DNA_w/601_, with the respective energies *E*_*left*_ and *E*_*right*_, are considerably decreased compared to tetrasomes loaded onto DNA_random_ (**Table II**). The heights of the energy barriers *ΔG\**_*left*_ and *ΔG\**_*right*_ are estimated from the rates *k*_*l->r*_ and *k*_*r->l*_, respectively. For tetrasomes on DNA_w/601_ relative to DNA_random_ in buffer A, *ΔG\**_*left*_ decreases strongly, whereas *ΔG\**_*right*_ is largely unchanged. In the presence of mono- and divalent salts in buffers B and C, tetrasomes on DNA_w/601_ become considerably longer-lived in the left-handed conformation (i.e. *ΔG\**_*left*_ is higher) compared to tetrasomes on DNA_w/601_ in buffer A, while the right-handed state (hence *ΔG\**_*right*_) remains essentially unaltered.

Further details of the data analysis have been described in Ref. (37). The significance of the similarity between the structural quantities obtained in the different conditions was assessed by respective unpaired two-sample t-tests (**Supplementary Tables SV-SVIII** and **SX**). The results for the dwell times obtained in the different conditions were checked for similarity by respective Wilcoxon rank-sum tests (**Supplementary Tables SXIII-SXV**). Finally, we note that the times measured in this study are an upper boundary due to the finite bead response time. The errors reported for the mean values obtained in this study correspond to 1 STD from the underlying distributions, unless indicated otherwise. The errors of calculated quantities were computed by error propagation.

### Construction of energy diagrams

To construct the schematic energy diagrams of the left- and right-handed states of tetrasomes (**Figure 3D, Supplementary Figure S8D**), we have computed the free energy difference between them and estimated the height of the barrier separating them. The free energy difference can be calculated from the ratio of the probabilities *p*_*left*_ = *p* and *p*_*right*_ = *1-p* for a tetrasome to be in the left- and right-handed conformations, respectively, using *ΔE*_*peak* *areas*_ = −k_B_Tln((*1-p*)/*p*) (see **Data Analysis** above, and **Supplementary Table SXI**). It can alternatively be computed from the ratio of the dwell times according to *ΔE*_*dwell-times*_ = −k_B_Tln(*τ*_*D,right*_/*τ*_*D,left*_). The obtained results are summarized in **Table II** and **Supplementary Table SXII**. While the results from both approaches agree within error, we consider the energy differences calculated from the dwell times as more reliable due to the larger sample size. To estimate the height of the energy barrier *ΔG\** relative to the left- and right-handed states of tetrasomes, we compute *ΔG* =* −k_B_Tln(*k*/*k*_*0*_), where the transition rates *k*_*l->r*_ and *k*_*r->l*_ equal the inverse of the dwell times *τ*_*D,left*_ and *τ*_*D,right*_, respectively, and the transition rate between the two states at zero force *k*_*0*_ is assumed to be symmetric and estimated at ∼10^7^ s^−1^ (37).

## RESULTS

### DNA sequence and buffer conditions do not affect the structural properties of tetrasomes

To assess whether high-affinity DNA sequences impact the formation and structure of tetrasomes, we loaded (H3.1-H4)_2_ tetramers onto individual DNA molecules containing a single centrally positioned high-affinity 601-sequence (DNA_w/601_) known for its strong nucleosome-positioning capability (50) with the help of the histone chaperone NAP1. Using FOMT (49), tetrasome assembly was recorded in real time by tracking the length *z* and linking number *Θ* of the DNA. In this setting, a single immobilized DNA molecule is tethered to a magnetic bead that is subject to constant force applied by a cylindrical permanent magnet, allowing its free rotation in the (*x,y*)-plane (**Figure 1B**). This force was set to 0.6-0.7 pN for direct comparison to our previous study of (H3.1-H4)_2_ tetrasomes assembled by histone chaperone NAP1 on DNA molecules of random sequence (DNA_random_) (35).

As in our previous studies with DNA_random_ (35-37), flushing in pre-incubated histone/NAP1-complexes (*n*_*exp,subset*_=9 out of total *N*_*exp,total*_=20) induced a simultaneous step-like change in both the length *z* and linking number *Θ* of DNA_w/601_, indicating the assembly of a single tetrasome (**Figure 1C**). The mean values of these changes (**Supplementary Table SIII**) are in excellent agreement with those previously obtained for NAP1-loaded (H3.1-H4)_2_ tetrasomes on DNA_random_ (35). Also in agreement with previous observations (35-37) is the lack of observable interactions between NAP1 chaperones and DNA molecules in the absence of histones (**Supplementary Figure S3**). To probe the effects of ambient conditions on tetrasome structure and dynamics, we performed the experiments in three different buffers employed in prior studies (35-37,51-53) that mainly differed in the concentrations of mono- and divalent salts (**Table I**) and obtained highly similar results (**Supplementary Table SIII**). We also flushed in (H3.1-H4)_2_ tetramers in the absence of NAP1 (*n*_*exp,subset*_=11). Thereby, DNA_w/601_ also exhibited a simultaneous step-like change in both measured quantities (**Supplementary Figure S4**). The mean values of the changes in length *z* and linking number *Θ* (**Supplementary Table SIII**) correspond excellently to those we obtained for (H3.1-H4)_2_ tetrasomes loaded by NAP1 onto DNA_w/601_, or onto DNA_random_ (35). The similarity of these results, which were furthermore independent of buffer conditions, indicate the formation of proper complexes in all cases. This was unexpected, since we never observed tetrasome formation in the absence of NAP1 in previous studies using random DNA sequences (35-37). Thus, the specific features of the high-affinity 601-sequence are capable of inducing spontaneous chaperone-independent assembly of histone tetramers.

Under the highly diluted conditions required for the controlled assembly of single tetrasomes (**Materials and Methods**), we also observed disassembly of tetrasomes in 40% (*n*_*exp,subset*_=8) of all experiments (*N*_*exp,total*_=20) within 3364 ± 765 s (mean ± 1 standard error of the mean (SEM); **Supplementary Figure S6**). The absolute mean values of the disassembly-associated changes in DNA length *z* and linking number *Θ* (**Supplementary Table SIV**) are in excellent agreement with those obtained upon assembly of tetrasomes (**Supplementary Table SIII**). Because disassembly occurred for both NAP1-loaded and spontaneously loaded tetrasomes in all buffer conditions, we can exclude a destabilizing effect of NAP1 and/or specific buffer conditions.

Given the similarity of the absolute changes in DNA length *z* and linking number *Θ* in all experiments in different buffer conditions (*N*_*exp,total*_=20) (as assessed by respective t-tests, **Supplementary Tables SV-SVIII**), all results were grouped together to yield improved statistics. The absolute changes in DNA length *z* and DNA linking number *Θ* upon assembly and disassembly of single tetrasomes (*N*_*counts,total*_=27) yielded mean values of *Δz*_*(dis)*ass_ = 23 ± 5 nm (*n*_*counts,subset*_=25) and *ΔΘ*_*(dis)ass*_ = 0.9 ± 0.2 turns (*n*_*counts,subset*_=26), respectively (**Figure 1D**). The distributions of the changes upon assembly or disassembly alone are shown in **Supplementary Figure S5A**,**B**, respectively. These results are in excellent agreement with values reported in other studies involving tetrasomes (25,30-37). Overall, our findings indicate that the high-affinity 601 sequence fully supports the assembly of (H3.1-H4)_2_ tetramers into proper tetrasome complexes, irrespective of the presence or absence of NAP1, or of buffer conditions.

### High-affinity tetrasomes exhibit spontaneous DNA handedness flipping

To investigate the time-dependent behavior of tetrasomes stably bound to DNA_w/601_, the FOMT measurements were continued for several hours following assembly. Like stably bound tetrasomes on DNA_random_ (35-37), high-affinity tetrasomes exhibited spontaneous dynamics in the handedness of their DNA wrapping (**Figure 2A**) irrespective of the buffer conditions. While the length *z* of the DNA molecules remained constant, the linking number *Θ* spontaneously fluctuated (or flipped) between two distinct mean values (**Figure 2B, Supplementary Figure S7**). Based on the negative and positive signs of these values, we deduce that they correspond to conformations of left- and right-handed DNA wrapping, respectively. As observed with tetrasomes on DNA_random_ (35-37), tetrasomes on DNA_w/601_ most frequently adopted the left-handed conformation. The difference in mean DNA linking number between the two observed states quantifies the induced change in tetrasome structure. The values of this quantity obtained under different experimental conditions, including tetrasomes loaded in the absence of NAP1, are summarized in **Supplementary Table SIX**. Due to the similarity of their values (as assessed by respective t-tests, **Supplementary Table SX**), the results from all experiments were pooled, which yielded a mean change in DNA linking number *Θ* upon flipping of *ΔΘ*_*flipping*_ = 1.6 ± 0.2 turns (*N*_*counts,total*_=23) (**Figure 2C**). These values are in excellent agreement with previous studies (35-37). Overall, our findings indicate that DNA sequence, the presence or absence of NAP1 during assembly, or buffer conditions do not affect the structural rearrangement of tetrasomes associated with handedness flipping.

### High-affinity DNA sequences increase the kinetics of tetrasome handedness flipping

We further examined the handedness dynamics of single tetrasomes in terms of the times spent in each of the two states of DNA wrapping (**Materials and Methods**). To specifically examine the role of DNA sequence, we focused these studies on NAP1-assembled tetrasomes, comparing the dynamics of NAP1-assembled (H3.1-H4)_2_ tetramers on either DNA_w/601_ or DNA_random_. The details of tetrasome handedness dynamics upon spontaneous, NAP1-independent assembly can be found in the **Supplementary Results and Discussion** and in **Supplementary Figure S8**. Our results indicate that the underlying DNA sequence significantly alters the kinetics of tetrasome handedness flipping (as assessed by respective Wilcoxon rank-sum tests, **Supplementary Tables SXIII-SXV**). This can be deduced from the observation that a tetrasome on DNA_w/601_ spent an average time of *τ*_*left*_ = 37 s in the left-handed state (in standard buffer A, **Figure 3A**, left), which is 4.9 ± 0.5 fold shorter than a tetrasome assembled onto DNA_random_ in the same buffer (*τ*_*left*_ = 183 s, **Table II**). The right-handed state was affected to a lesser extent, displaying a lifetime *τ*_*right*_ = 28 s (in standard buffer A, **Figure 3A**, right), which is 2.0 ± 0.3 times longer than a tetrasome on DNA_random_ in the same buffer (*τ*_*right*_ = 14 s, **Table II**). These results clearly show that tetrasomes on high-affinity DNA sequences have a substantially enhanced tendency to flip from the left- to the right-handed state compared to tetrasomes on random DNA sequences (**Supplementary Table SXIII**). Since the experimental conditions were otherwise identical, these effects must arise from the 601-sequence in DNA_w/601_.

We can translate the information from dwell-time measurements into schematic energy diagrams depicting the left- and right-handed states of tetrasomes (**Materials and Methods**; **Table II** and **Figure 3D**). Notably, the free energy differences *ΔE* between the left- and right-handed conformations of tetrasomes are reduced ∼9-fold on DNA_w/601_ (*ΔE* = 0.3 ± 0.1 k_B_T) relative to DNA_random_ (*ΔE* = 2.6 ± 0.1 k_B_T), reflecting the shift of the equilibrium occupancy towards the right-handed conformation for NAP1-loaded tetrasomes on DNA_w/601_. This results predominantly from the smaller estimated energy difference between the left-handed state and the transition state (at the top of the barrier) for tetrasomes loaded onto DNA_w/601_ (*ΔG\**_*left*_∼19.7 k_B_T) relative to DNA_random_ (*ΔG\**_*left*_∼21.3 k_B_T).

We next examined the flipping dynamics of tetrasomes on DNA_w/601_ in two different buffer conditions. In buffer B (containing a two-fold increased concentration of monovalent salt in the absence of crowding agents, **Table I**), tetrasomes dwelled in the left-handed state for *τ*_*left*_ = 62 s (**Figure 3B**, left), 1.7 ± 0.1 times longer than in standard buffer A (*τ*_*left*_ = 37 s, **Table II**; **Figure 3A**, left). The average lifetimes in the right-handed conformation were affected only slightly by the change to buffer B (**Figure 3B**, right; **Table II, Supplementary Tables SXIV** and **SXV**). In buffer C, which contained additional magnesium chloride relative to buffer B (**Table I**), a tetrasome dwelled in the left-handed conformation for an even longer time of *τ*_*lef*t_ = 141 s (**Figure 3C**, top), which is 2.3 ± 0.2-fold longer than in buffer B (*τ*_*left*_ = 62 s, **Table II**; **Figure 3B**, left). Again, the mean dwell time in the right-handed state showed essentially no change (**Figure 3C**, bottom; **Figure 3B**, right).

These findings can also be transferred into schematic energy diagrams (**Materials and Methods**; **Figure 3D**). Because the predominant effect of increased monovalent salt concentration in buffer B for tetrasomes on DNA_w/601_ is an increase in the lifetime of the left-handed conformation, the free energy difference between the two states increases to *ΔE* = 0.6 ± 0.1 k_B_T (compared to *ΔE* = 0.3 ± 0.1 k_B_T in buffer A; **Table II**). The estimated barrier height relative to the left-handed state is altered to *ΔG\**_*left*_∼20.2 k_B_T, while the barrier height relative to the right-handed state remains essentially unaffected. Similarly, because the higher concentration of divalent ions in buffer C caused a further extension of the lifetime of the left-handed conformation, the free energy difference between the two states correspondingly increases to *ΔE* = 1.5 ± 0.1 k_B_T (**Table II**). Thus, the estimated barrier height relative to the left-handed state for tetrasomes on DNA_w/601_ is larger in buffer C (*ΔG\**_*left*_∼21.1 k_B_T) than in buffer B (*ΔG\**_*left*_∼20.2 k_B_T,), while the barrier height relative to the right-handed state remains essentially unaffected. In summary, higher concentrations of monovalent salt in the absence of crowding agents as well as divalent salts attenuate the handedness flipping dynamics of tetrasomes assembled on the high-affinity Widom 601-sequence by increasing the dwell time in the left-handed state and, thus, reducing the tendency for tetrasomes to flip to the right-handed state.

## DISCUSSION

A strong component determining the positioning and stability of nucleosomes throughout chromatin is the DNA sequence (63). For instance, nucleosome maps obtained from chromatin assembled *in vitro* on purified yeast genomic DNA showed remarkable similarity to nucleosome maps from native chromatin (64). The observation that specific periodic patterns of AA, TT, TA or GC dinucleotides favour DNA wrapping around the histone core led to the development of algorithms to predict nucleosome positions in different organisms (42,65,66). High-affinity nucleosome binding sequences have been found to be enriched at the transcriptional start site, in intragenic locations or over genes subject to strong regulation (42). In the present study, we have investigated to which extent such high-affinity sites affect the structure and dynamics of subnucleosomal (H3-H4)_2_ tetrasomes. Our previous single-molecule experiments revealed that the left-handed wrapping of DNA around the histone octamer is predominantly static, while the absence of H2A-H2B dimers results in highly dynamic flipping of DNA handedness between the left- and a right-handed state (35-37). However, these studies were carried out on tetrasomes assembled onto random DNA sequences, and it remained unclear whether high-affinity sequences, exemplified by the well-characterized Widom 601-sequence (50), would have an impact on the dynamics of tetrasomes. Our assessment of the assembly and structural features of (H3-H4)_2_ tetrasomes on DNA including the 601-sequence revealed that the key characteristics of assembled tetrasomes were very similar to those assembled onto random DNA (35-37). Interestingly, however, we observed that the 601 sequence also supported spontaneous, histone chaperone-independent formation of tetrasomes. As this property strongly distinguishes DNA including the high-affinity 601-sequence from DNA of random sequence, this suggests that tetrasome assembly occurs at the 601-sequence under these conditions. In the following, we assume that a similar sequence-specificity holds for the chaperoned assembly of tetrasomes onto DNA including the 601-sequence.

In contrast to the overall structural similarity between tetrasomes assembled onto high-affinity versus random DNA sequences, our analyses revealed a striking difference with respect to handedness flipping dynamics. The presence of the 601-sequence considerably decreased the lifetime of the canonical left-handed state of high-affinity tetrasomes, corresponding to a reduction in the free energy *ΔE* between the left- and right-handed conformations, which results in enhanced flipping frequency.

These results may at first sight seem surprising given the high propensity of the 601-sequence to support nucleosome/tetrasome formation relative to other DNA sequences (67), which presumably derives from a combination of enhanced binding to histones and increased bendability or twistability of the DNA (38,50). Indeed, our observation that histone tetramers assemble spontaneously onto DNA molecules with a 601-sequence, but not onto DNA molecules of random sequence, emphasizes the strong affinity of the sequence for histones. Yet what could be the reason for the increased flipping dynamics of high-affinity tetrasomes? A likely explanation of these findings could be that the increased binding stability of tetrasomes to DNA reduces the influence of thermal fluctuations on the disruption of histone-DNA contacts (by analogy to nucleosome ‘breathing’ (11-14)), and instead transmits thermal effects to the H3-H3 interface. This could promote the rotation of the H3-H4 dimers relative each other that is proposed to underlie handedness flipping (30-37). In this model, high-affinity sequences might not only promote energetically favorable histone-DNA interactions, but may also endow tetrasomes with enhanced ability to accommodate torsional stress by flipping between left- and right-handed conformation. This feature may be particularly important during transcription, when tetrasomes can arise from nucleosomes by loss or removal of H2A/H2B dimers, and when torque can be generated by elongating RNA polymerase II (68-70). Although most transcription-linked torsional stress is removed by topoisomerases (71,72), high-affinity tetrasomes may provide extra protection against complete histone ejection at crucial locations in the genome by their inherently stronger ability to absorb this stress.

In the presence of higher concentrations of mono- and divalent salt and an absence of synthetic crowding agents, the left-handed states of NAP1-loaded tetrasomes on DNA molecules with a 601-sequence become longer lived. Hence, these conditions decrease the tendency of tetrasomes to adopt the right-handed conformation. We consider a direct influence of the synthetic crowding agents PEG and PVA on tetrasome dynamics to be unlikely, as studies show that the influence of PEG (73) or PVA (74) on the kinetics of biological systems is negligible at the low concentrations (below 1%) employed in our standard buffer. Thus, these changes must arise from the higher concentration of salts, which will reduce the strength of electrostatic interactions through screening. Even a two-fold increase in salt concentration as employed here can induce small but significant changes in the properties of DNA (e.g. elasticity (75-77), torsional stiffness (78), and supercoiling (79)) and proteins (80), and in the case of histones, reduce their affinity for DNA (81). In agreement with the model proposed above, such a generic reduction in affinity could account for the observed increase of the lifetime of the left-handed state for tetrasomes on DNA molecules with a 601-sequence, because it would shift the balance between thermally induced opening of DNA-histone contacts and dimer rotation at the H3-H3 interface towards the former. Upon further addition of divalent salt, which increases electrostatic screening to an even larger extent, the dwell time in the left-handed state of NAP1-loaded tetrasomes on DNA molecules with a 601-sequence becomes more pronounced, as one might expect by analogy with the above. These findings also illustrate that the interpretation of results obtained using nucleosome-positioning sequences must take into account the specific buffer conditions employed.

In summary, we provide conceptual evidence that DNA sequence impacts tetrasome handedness flipping dynamics, which may contribute to the mitigation of torsional stress generated by molecular motors without accompanying eviction of histones.

## AUTHOR CONTRIBUTIONS

O.O., A.L., and N.H.D. designed the research; O.O. performed the research and analyzed the data; O.O., A.L., and N.H.D. discussed the results and wrote the paper.

## ACKNOWLEDGMENTS

The authors thank Artur Kaczmarczyk for insightful discussions, the laboratory of John van Noort for providing the pGEM3Z-mono601 plasmid and several stock solutions, Theo van Laar for preparing the DNA constructs, and Jacob Kerssemakers for providing the improved step-finder algorithm. The authors also acknowledge financial support from the funding agencies mentioned below.

## SUPPLEMENTARY DATA

A Supplementary Information file accompanies this manuscript.

