## Supplementary Information for "DNA sequence is a major determinant of tetrasome dynamics"

O. Ordu, A. Lusser, and N. H. Dekker

### SUPPLEMENTARY RESULTS AND DISCUSSION

**The dwell times in the left-handed state, and the free energy difference between the left- and right-handed states, are increased for spontaneously loaded tetrasomes compared to NAP1-loaded tetrasomes on $DNA_{w/601}$**

We have measured the dwell times of tetrasomes assembled in the absence of NAP1 ('spontaneously loaded tetrasomes') under the standard conditions in buffer A (**Table SXII** and **Figure S8A)**. These displayed an average dwell time in the left-handed conformation of $\tau_{left}$ = 93 s (**Table SXII** and **Figure S8A**, left), 2.5 ± 0.2 times longer than that for NAP1-loaded tetrasomes on $DNA_{w/601}$ ($\tau_{left}$ = 37 s, **Table II** and **Figure 3A**, left, in the **main text**). Conversely, the dwell time in the right-handed state was only slightly increased to $\tau_{right}$ = 34 s (**Table SXII** and **Figure S8A**, right), from $\tau_{right}$ = 28 s for NAP1-loaded tetrasomes on $DNA_{w/601}$ (**Table II** and **Figure 3A**, right, in the **main text**). While the dwell time in the left-handed state is thus considerably longer-lived for spontaneously loaded tetrasomes than for NAP1-loaded tetrasomes on $DNA_{w/601}$, it still remains remarkably shorter than that of NAP1-loaded tetrasomes on $DNA_{random}$ ($\tau_{left}$ = 183 s**, Table II** in the **main text**, and **Table SXIII**) by 2.0 ± 0.2 fold. In the right-handed state, the dwell time for spontaneously loaded tetrasomes considerably exceeds that of NAP1-loaded tetrasomes on $DNA_{random}$ ($\tau_{right}$ = 14 s, **Table II** in the **main text**, and **Table SXIII**) by 2.4 ± 0.3 fold.

In the main text, we have shown that the free energy difference (**Materials and Methods** in the **main text**) between the left- and right-handed conformations is larger for tetrasomes loaded by NAP1 on $DNA_{random}$ ($\Delta E$ = 2.6 ± 0.1 $k_BT$) than for tetrasomes loaded by NAP1 on $DNA_{w/601}$ ($\Delta E$ = 0.3 ± 0.1 $k_BT$) (**Table II** and **Figure 3D** in the **main text**). For spontaneously loaded tetrasomes, the free energy difference ($\Delta E$ = 1.0 ± 0.1 $k_BT$) lies in between these values (**Table SXII** and **Figure S8D**). We have also shown that the estimated barrier height (**Materials and Methods** in the **main text**) separating the left-handed state from the transition state is higher for NAP1-loaded tetrasomes on $DNA_{random}$ ($\Delta G^*_{left}$~21.3 $k_BT$) than for NAP1-loaded tetrasomes on $DNA_{w/601}$ ($\Delta G^*_{left}$~19.7 $k_BT$). The barrier height for spontaneously loaded tetrasomes, $\Delta G^*_{left}$~20.7 $k_BT$, lies in between these values.

We have also measured the dwell times of spontaneously loaded tetrasomes in buffers B and C (**Table SXII** and **Figure S8B**,**C**). In buffer B, spontaneously loaded tetrasomes displayed increased dwell times in the left-handed state ($\tau_{left}$ = 175 s, **Figure S8B**, left, 1.9 ± 0.1 times longer than in buffer A where $\tau_{left}$ = 93 s, **Figure S8A**, left; **Table SXII**), whereas the average lifetimes in the right-handed conformation were only slightly affected (**Tables SXII**, **SXIV**, **SXV**, and **Figures S8A,B**, right). Hence, the average dwell time of spontaneously loaded tetrasomes in the left-handed state remained considerably shorter than that of NAP1-loaded tetrasomes by a factor of 2.8 ± 0.2 (**Table II** and **Figure 3B**, left, in the **main text**; **Tables SXII**, **SXIV**, **SXV**, and **Figure S8B**, left). In buffer C, spontaneously loaded tetrasomes were observed to dwell in the left-handed state for shorter times ($\tau_{left}$ = 93 s, **Figure S8C**, top; 1.9 ± 0.1 times shorter than in buffer B where $\tau_{left}$ = 175 s, **Figure S8B**, left; **Table SXII**). In this buffer, a similarly sized decrease in the average dwell time was observed in the right-handed conformation ($\tau_{right}$ = 19 s, **Figure S8C**, bottom; 1.4 ± 0.1-fold shorter than in buffer B where $\tau_{left}$ = 30 s, **Figure S8B**, right; **Table SXII**). As a result, in buffer C, the average dwell time of a spontaneously loaded tetrasome in the left-handed state decreased considerably compared to that of a NAP1-loaded tetrasome, by a factor of 1.5 ± 0.1; for the right-handed conformation, a similar factor of 1.6 ± 0.1 was found (**Table II** and **Figure 3C** in the **main text**; **Tables SXII, SXIV, SXV** and **Figure S8C**).

As shown in the schematic energy diagrams (**Materials and Methods** in the **main text**) in **Figure S8D**, for spontaneously loaded tetrasomes the free energy difference between the left- and right-handed states increased to $\Delta E$ = 1.9 ± 0.1 $k_BT$ in buffer B (compared to $\Delta E$ = 1.0 ± 0.1 $k_BT$ in buffer A, **Table SXII**). The estimated barrier height relative to the left-handed state remained larger for spontaneously loaded tetrasomes ($\Delta G^*_{left}$~21.3 $k_BT$) than for NAP1-loaded tetrasomes on $DNA_{w/601}$ ($\Delta G^*_{left}$~20.2 $k_BT$), while the barrier height relative to the right-handed state remained essentially unaltered. Because a change to buffer C left the free energy difference between the two states largely unaffected ($\Delta E$ = 1.6 ± 0.1 $k_BT$ versus $\Delta E$ = 1.9 ± 0.1 $k_BT$, **Table SXII** and **Figure S8D**), in this buffer the free energy difference between the two states was similar to that of NAP1-loaded tetrasomes on $DNA_{w/601}$ ($\Delta E$ = 1.5 ± 0.1 $k_BT$, **Table II** in the **main text**).

These results indicate that the flipping kinetics of tetrasomes differ depending on whether a histone chaperone, such as NAP1, was present during their assembly. This effect is, however, less pronounced than the effect of DNA sequence highlighted in the main text. Possibly, the chaperoning provided by histone chaperones such as NAP1 leads to a more stable assembly of tetrasomes at the 601 sequence. Alternatively, the spontaneous association of $(H3\text{-}H4)_2$ tetramers onto high-affinity DNA, while possible, could lead to subtle differences in tetrasome conformation with slightly weaker histone-DNA contacts. This may interfere with the transmission of thermal forces to the H3-H3 interface, which in turn may account for the reduced handedness flipping dynamics of spontaneously loaded tetrasomes (see **main text** for an in-depth discussion of the impact of DNA sequence on tetrasome flipping dynamics).

### SUPPLEMENTARY TABLES

**Supplementary Table SI. The sequences of the primers used to make the main DNA fragments.** The specific overhangs at the *Bsa*I sites are indicated in lowercase.

| Primers | Primer Sequences |
|---|---|
| **1** | 5'-CCATCTTGGTCTCCtcaaatcctgttaccagtggctgctgcc |
| **2** | 5'-CCATCTTGGTCTCCtaggtgttgagatccagttcgatgtaacc |
| **3** | 5'-CCATCTTGGTCTCCttgaCGCTCTAGAACTAGTGGATCCCCC |
| **4** | 5'-CCATCTTGGTCTCCcctaCGCTCTAGAACTAGTGGATCCCCC |
| **5** | 5'-GACCGAGATAGGGTTGAGTG |

**Supplementary Table SII. Concentrations and volumes of the DNA and bead solutions used in this work.** The remaining details of the flow cell and sample preparation have been described in Ref. (1).

| Sample | Concentrations and volumes |
|---|---|
| **DNA molecules (Figure S1)** | Stock concentration: $c_{stock}$ = 45 ng/µl;<br>For tethering: 0.2 µl of 1:300 dilution from stock solution |
| **superparamagnetic beads (0.5-µm diameter, Ademtech, Pessac, France)** | For tethering: 0.2 µl or 0.3 µl from stock solution |

**Supplementary Table SIII. Summary of the assembly events.** Sizes of the step-like change in DNA length $\Delta z_{ass}$ (in nm) and $\Delta\Theta_{ass}$ (in turns) upon the assembly of a single tetrasome in the different buffer conditions. The compositions of the buffers are provided in **Table I** of the **main text**.

| Buffers | $\Delta z_{ass}$ (nm) | $\Delta\theta_{ass}$ (turns) | mean $\Delta z_{ass}$ (± STD, nm) | mean $\Delta\theta_{ass}$ (± STD, turns) |
|---|---|---|---|---|
| **Buffer A** | *-21* | *-1.0* | | |
| ***green: with NAP1*** | *-23* | *-1.2* | | |
| | *-25* | *-0.7* | *-23 (± 2)* | *-1.0 (± 0.3)* |
| **blue: without NAP1** | -23 | -0.8 | | |
| | -10 | -0.8 | | |
| | -23 | -1.3 | | |
| | -28 | -0.5 | -25 (± 3)* | -1.0 (± 0.3)** |
| ***overall means buffer A*** | | | ***-24 (± 3)**** | ***-1.0 (± 0.2)***** |
| **Buffer B** | *-16* | *-1.3* | | |
| | *-16* | *-1.0* | | |
| | *-24* | *-1.0* | | |
| | *-16* | *-0.9* | | |
| | *-33* | *-0.7* | *-21 (± 7)* | *-1.0 (± 0.2)* |
| | -23 | -0.9 | - | - |
| ***overall means buffer B*** | | | ***-21 (± 7)*** | ***-1.0 (± 0.2)*** |
| **Buffer C** | -24 | -0.6 | | |
| | -27 | -0.7 | | |
| | -19 | -0.8 | | |
| | -16 | -1.2 | -22 (± 5) | -0.8 (± 0.2) |
| **TOTAL MEANS** | | | **-22 (± 5)*** | **-0.9 (± 0.2)**** |

*excluding the value ($\Delta z_{ass}$ = -10 nm) beyond the mean resolution in $z$ ($z_{res}$ = ±16 nm)

**excluding the value $\Delta\Theta_{ass}$ = -0.5 (-0.46) turns beyond the mean resolution in $\Theta$ ($\Theta_{res}$ = ±0.5 (0.47) turns)

**Supplementary Table SIV. Summary of the disassembly events.** Sizes of the step-like change in DNA length $\Delta z_{disass}$ (in nm) and $\Delta\Theta_{disass}$ (in turns) upon the disassembly of a single tetrasome in the different buffer conditions. The compositions of the buffers are provided in **Table I** of the **main text**.

| Buffers | $\Delta z_{disass}$ (nm) | $\Delta\theta_{disass}$ (turns) | mean $\Delta z_{disass}$ (± STD, nm) | mean $\Delta\theta_{disass}$ (± STD, turns) |
|---|---|---|---|---|
| **Buffer A** | ***31*** | ***0.7*** | | |
| ***green: NAP1-loaded*** | ***19*** | ***0.5*** | ***25 (± 8)*** | ***0.6 (± 0.1)*** |
| **blue: without NAP1** | **23** | **0.6** | | |
| | **20** | **0.9** | | |
| | **25** | **-1.1** | **23 (± 3)** | **0.9 (± 0.2)**** |
| ***overall means in buffer A*** | | | ***24 (± 5)*** | ***0.8 (± 0.2)***** |
| **Buffer B** | ***11*** | ***0.9*** | | |
| | ***25*** | ***0.7*** | | |
| | ***25*** | ***1.2*** | ***25 (± 0)**** | ***0.9 (± 0.2)*** |
| | **33** | **0.7** | **-** | **-** |
| ***overall means in buffer B*** | | | ***28 (± 5)**** | ***0.9 (± 0.2)*** |
| **Buffer C** | **22** | **0.6** | **-** | **-** |
| **TOTAL MEANS** | | | **25 (± 5)*** | **0.8 (± 0.2)**** |

*excluding the value ($\Delta z_{disass}$ = 11 nm) beyond the mean resolution in $z_{res}$ = ±16 nm

**considering the absolute value from the single right-handed disassembly observed ($\Delta\Theta_{disass}$ = -1.1 turns)

**Supplementary Table SV. Results of the unpaired two-sample t-tests on the changes in DNA length *Δz* upon tetrasome assembly or disassembly.** An h-value of 0 (1) indicates that the data sets (do not) originate from independent, normally distributed random samples with equal means and equal but unknown variances at 5% significance level, i.e. with 95% confidence. The p-value essentially corresponds to the probability of the similarity between the two data sets.

**DISASSEMBLY**

| Buffers + or *-NAP1* | Buffer A *+NAP1* | Buffer B *+NAP1* | Buffer C *+NAP1** | Buffer A *-NAP1* | Buffer B *-NAP1*** | Buffer C *-NAP1(**)* |
|---|---|---|---|---|---|---|
| Buffer A *+NAP1* | ---- | h = 0<br>p = 0.98 | ---- | h = 0<br>p = 0.65 | h = 0<br>p = 0.59 | h = 0<br>p = 0.79 |
| Buffer B *+NAP1* | *h = 0<br>p = 0.68* | ---- | ---- | h = 0<br>p = 0.34 | h = 1<br>p = 0.02 | h = 0<br>p = 0.05 |
| Buffer C *+NAP1** | ---- | ---- | ---- | ---- | ---- | ---- |
| Buffer A *-NAP1* | *h = 0<br>p = 0.57* | *h = 0<br>p = 0.49* | ---- | ---- | h = 0<br>p = 0.08 | h = 0<br>p = 0.75 |
| Buffer B *-NAP1*** | *h = 0<br>p = 0.98* | *h = 0<br>p = 0.83* | ---- | *h = 0<br>p = 0.72* | ---- | h = NaN<br>p = NaN |
| Buffer C *-NAP1(**)* | *h = 0<br>p = 0.59* | *h = 0<br>p = 0.94* | ---- | *h = 0<br>p = 0.37* | *h = 0<br>p = 0.78* | ---- |

***ASSEMBLY***

*neither assembly nor disassembly data available; **data sets contain one value;
(**)disassembly data set contains one value

**Supplementary Table SVI. Results of the unpaired two-sample t-tests on the changes in DNA length *Δz* upon tetrasome assembly and disassembly.** An h-value of 0 indicates that the data sets originate from independent, normally distributed random samples with equal means and equal but unknown variances at 5% significance level, i.e. with 95% confidence. The p-value essentially corresponds to the probability of the similarity between the two data sets.

| ASSEMBLY \ DISASSEMBLY | | | | | | |
|---|---|---|---|---|---|---|
| **Buffers + or -*NAP1*** | **Buffer A +*NAP1*** | **Buffer B +*NAP1*** | **Buffer C +*NAP1**** | **Buffer A -*NAP1*** | **Buffer B -*NAP1***** | **Buffer C -*NAP1***** |
| **Buffer A +*NAP1*** | h = 0<br>p = 0.70 | h = 0<br>p = 0.37 | ---- | h = 0<br>p = 0.84 | h = 0<br>p = 0.07 | h = 0<br>p = 0.62 |
| **Buffer B +*NAP1*** | h = 0<br>p = 0.56 | h = 0<br>p = 0.52 | ---- | h = 0<br>p = 0.74 | h = 0<br>p = 0.23 | h = 0<br>p = 0.96 |
| **Buffer C +*NAP1**** | ---- | ---- | ---- | ---- | ---- | ---- |
| **Buffer A -*NAP1*** | h = 0<br>p = 0.90 | h = 0<br>p = 0.85 | ---- | h = 0<br>p = 0.48 | h = 0<br>p = 0.14 | h = 0<br>p = 0.48 |
| **Buffer B -*NAP1***** | h = 0<br>p = 0.87 | h = 0<br>p = 0.09 | ---- | h = 0<br>p = 0.92 | h = NaN<br>p = NaN | h = NaN<br>p = NaN |
| **Buffer C -*NAP1*** | h = 0<br>p = 0.50 | h = 0<br>p = 0.36 | ---- | h = 0<br>p = 0.70 | h = 0<br>p = 0.11 | h = 0<br>p = 0.98 |

*no data available; **data sets contain one value

**Supplementary Table SVII. Results of the unpaired two-sample t-tests on the changes in DNA linking number *ΔΘ* upon tetrasome assembly or disassembly.** An h-value of 0 indicates that the data sets originate from independent, normally distributed random samples with equal means and equal but unknown variances at 5% significance level, i.e. with 95% confidence. The p-value essentially corresponds to the probability of the similarity between the two data sets.

| ASSEMBLY \ DISASSEMBLY | | | | | | |
|---|---|---|---|---|---|---|
| **Buffers + or -*NAP1*** | **Buffer A +*NAP1*** | **Buffer B +*NAP1*** | **Buffer C +*NAP1**** | **Buffer A -*NAP1*** | **Buffer B -*NAP1***** | **Buffer C -*NAP1*$^{(**)}$** |
| **Buffer A +*NAP1*** | ---- | h = 0<br>p = 0.18 | ---- | h = 0<br>p = 0.31 | h = 0<br>p = 0.72 | h = 0<br>p = 0.97 |
| **Buffer B +*NAP1*** | *h = 0*<br>*p = 0.98* | ---- | ---- | h = 0<br>p = 0.75 | h = 0<br>p = 0.45 | h = 0<br>p = 0.35 |
| **Buffer C +*NAP1**** | ---- | ---- | ---- | ---- | ---- | ---- |
| **Buffer A -*NAP1*** | *h = 0*<br>*p = 0.94* | *h = 0*<br>*p = 0.94* | ---- | ---- | h = 0<br>p = 0.63 | h = 0<br>p = 0.50 |
| **Buffer B -*NAP1***** | *h = 0*<br>*p = 0.71* | *h = 0*<br>*p = 0.62* | ---- | *h = 0*<br>*p = 0.76* | ---- | h = NaN<br>p = NaN |
| **Buffer C -*NAP1*$^{(**)}$** | *h = 0*<br>*p = 0.38* | *h = 0*<br>*p = 0.27* | ---- | *h = 0*<br>*p = 0.44* | *h = 0*<br>*p = 0.85* | ---- |

DISASSEMBLY (upper triangle) / ASSEMBLY (lower triangle)

*neither assembly nor disassembly data available; **data sets contain one value; $^{(**)}$disassembly data set contains one value

**Supplementary Table SVIII. Results of the unpaired two-sample t-tests on the changes in DNA linking number *ΔΘ* upon tetrasome assembly and disassembly.** An h-value of 0 indicates that the data sets originate from independent, normally distributed random samples with equal means and equal but unknown variances at 5% significance level, i.e. with 95% confidence. The p-value essentially corresponds to the probability of the similarity between the two data sets.

| | | DISASSEMBLY | | | | | |
|---|---|---|---|---|---|---|---|
| | **Buffers + or *-NAP1*** | **Buffer A *+NAP1*** | **Buffer B *+NAP1*** | **Buffer C *+NAP1**** | **Buffer A *-NAP1*** | **Buffer B *-NAP1***** | **Buffer C *-NAP1***** |
| ASSEMBLY | **Buffer A *+NAP1*** | h = 0<br>p = 0.19 | h = 0<br>p = 0.79 | ---- | h = 0<br>p = 0.60 | h = 0<br>p = 0.46 | h = 0<br>p = 0.37 |
|  | **Buffer B *+NAP1*** | h = 0<br>p = 0.09 | h = 0<br>p = 0.76 | ---- | h = 0<br>p = 0.52 | h = 0<br>p = 0.30 | h = 0<br>p = 0.21 |
|  | **Buffer C *+NAP1**** | ---- | ---- | ---- | ---- | ---- | ---- |
|  | **Buffer A *-NAP1*** | h = 0<br>p = 0.22 | h = 0<br>p = 0.87 | ---- | h = 0<br>p = 0.67 | h = 0<br>p = 0.51 | h = 0<br>p = 0.42 |
|  | **Buffer B *-NAP1***** | h = 0<br>p = 0.40 | h = 0<br>p = 0.78 | ---- | h = 0<br>p = 0.97 | h = NaN<br>p = NaN | h = NaN<br>p = NaN |
|  | **Buffer C *-NAP1*** | h = 0<br>p = 0.42 | h = 0<br>p = 0.49 | ---- | h = 0<br>p = 0.73 | h = 0<br>p = 0.75 | h = 0<br>p = 0.59 |

*no data available; **data sets contain one value

**Supplementary Table SIX. Summary of the flipping events.** Sizes of the step-like changes in DNA linking number $\Delta\Theta_{flipping}$ (in turns) upon the flipping of an assembled tetrasome in different buffer conditions. The buffer compositions are provided in **Table I** of the **main text**.

| **Buffers** | **$\Delta\theta_{flipping}$ (turns)** ***with NAP1*** | **$\Delta\theta_{flipping}$ (turns)** ***without NAP1*** | **mean $\Delta\theta_{flipping}$ (±STD, turns)** |
|---|---|---|---|
| **Buffer A** | ***1.8*** | **1.6** | |
| | ***1.0*** | **1.6** | |
| | ***1.8*** | **1.7** | |
| | **-** | **1.3** | |
| | **-** | **1.3** | |
| ***overall means in buffer A*** | ***1.5 (± 0.4)*** | **1.5 (± 0.2)** | ***1.5 (± 0.3)*** |
| **Buffer B** | ***1.6*** | **1.0** | |
| | ***1.8*** | **1.7** | |
| | ***1.7*** | **1.7** | |
| | ***1.7*** | **-** | |
| | ***1.3*** | **-** | |
| | ***1.2*** | **-** | |
| | ***1.7*** | **-** | |
| ***overall means in buffer B*** | ***1.6 (± 0.2)*** | **1.5 (± 0.4)** | ***1.5 (± 0.3)*** |
| **Buffer C** | ***1.7*** | **1.7** | |
| | **-** | **1.7** | |
| | **-** | **1.7** | |
| | **-** | **1.7** | |
| ***overall means in buffer B*** | **-** | **1.7 (± 0.0)** | ***1.7 (± 0.0)*** |
| **TOTAL MEAN** | | | **1.6 (± 0.2)** |

**Supplementary Table SX. Results of the unpaired two-sample t-tests on the changes in DNA linking number $\Delta\Theta_{flipping}$ upon tetrasome flipping.** An h-value of 0 indicates that the data sets originate from independent, normally distributed random samples with equal means and equal but unknown variances at 5% significance level, i.e. with 95% confidence. The p-value essentially corresponds to the probability of similarity between the two data sets.

| Buffers + or *-NAP1* | Buffer A *+NAP1* | Buffer B *+NAP1* | Buffer C *+NAP1** | Buffer A *-NAP1* | Buffer B *-NAP1* | Buffer C *-NAP1* |
|---|---|---|---|---|---|---|
| Buffer A *+NAP1* | --- | h = 0<br>p = 0.79 | h = 0<br>p = 0.73 | h = 0<br>p = 0.97 | h = 0<br>p = 0.89 | h = 0<br>p = 0.42 |
| Buffer B *+NAP1* | | --- | h = 0<br>p = 0.57 | h = 0<br>p = 0.62 | h = 0<br>p = 0.60 | h = 0<br>p = 0.29 |
| Buffer C *+NAP1** | | | --- | h = 0<br>p = 0.31 | h = 0<br>p = 0.63 | h = 0<br>p = 0.47 |
| Buffer A *-NAP1* | | | | --- | h = 0<br>p = 0.83 | h = 0<br>p = 0.05 |
| Buffer B *-NAP1* | | | | | --- | h = 0<br>p = 0.26 |

*data sets contain one value

**Supplementary Table SXI. Probabilities for a tetrasome on a DNA molecule with a 601-sequence to occupy the left- or right-handed state in the different buffer conditions.** The respective probabilities $p_{left} = p$ and $p_{right} = 1 - p$ were obtained from the relative ratio between the peak areas of the two Gaussian functions fitted to the linking number data (**Materials and Methods** and **Figure 2B** in the **main text**, and **Figure S7**).

| Buffers | *p* *with NAP1* | *1-p* *with NAP1* | *p* without NAP1 | *1-p* without NAP1 |
|---|---|---|---|---|
| Buffer A | *0.86* | *0.14* | 0.72 | 0.28 |
| | *0.51* | *0.49* | 0.80 | 0.20 |
| | *0.73* | *0.27* | 0.73 | 0.27 |
| | - | - | 0.80 | 0.20 |
| | - | - | 0.71 | 0.29 |
| *overall means in buffer A* | *0.70 (± 0.18)* | *0.30 (± 0.18)* | 0.75 (± 0.05) | 0.25 (± 0.05) |
| Buffer B | *0.83* | *0.17* | 0.90 | 0.10 |
| | *0.92* | *0.08* | 0.86 | 0.14 |
| | *0.83* | *0.17* | 0.90 | 0.10 |
| | *0.85* | *0.15* | - | - |
| | *0.46* | *0.54* | - | - |
| | *0.90* | *0.10* | - | - |
| | *0.81* | *0.19* | | |
| *overall means in buffer B* | *0.80 (± 0.15)* | *0.20 (± 0.15)* | 0.89 (± 0.02) | 0.11 (± 0.02) |
| Buffer C | *0.86* | *0.14* | 0.90 | 0.10 |
| | - | - | 0.85 | 0.15 |
| | - | - | 0.74 | 0.26 |
| | - | - | 0.85 | 0.15 |
| *overall means in buffer B* | - | - | 0.83 (± 0.07) | 0.17 (± 0.07) |

**Supplementary Table SXII. Summary of quantified properties for spontaneously loaded tetrasomes on DNA with a 601-sequence.** The times $\tau_{D,left}$ and $\tau_{D,right}$ that a tetrasomes loaded onto a $DNA_{w/601}$ molecule in the absence of NAP1 ('spontaneously loaded tetrasome') dwelled in the left- and right-handed states, respectively. Buffers employed are indicated within brackets. The underlying analysis is described in **Materials and Methods** in the **main text**, and detailed in Ref. (1).

| time average in filtering (s) | $\tau_{D,left}$ (s) [buffer type] | $\tau_{D,right}$ (s) [buffer type] | $\tau_{D,left}/\tau_{D,right}$ | $\Delta E_{dwell-times}$ ($k_BT$) | $\Delta E_{peak-areas}$ ($k_BT$) |
|---|---|---|---|---|---|
| **3.3 (N=330)** | 93 (+7/-6)** [A]<br>175 (+11/-10)** [B]<br>93 (±4)** [C] | 34 (+3/-2)** [A]<br>27 (±2)** [B]<br>19 (±1)** [C] | 3 ± 0* [A]<br>6 ± 0* [B]<br>5 ± 0* [C] | 1.0 ± 0.1* [A]<br>1.9 ± 0.1* [B]<br>1.6 ± 0.1* [C] | 1.1 ± 0.2* [A]<br>2.1 ± 0.2* [B]<br>1.6 ± 0.4* [C] |
| **19.1 (N=1910)** | 164 (+18/-15)** [A]<br>344 (+33/-28)** [B]<br>194 (+12/-11)** [C] | 58 (+6/-5)** [A]<br>50 (+5/-4)** [B]<br>36 (±2)** [C] | 3 ± 0* [A]<br>7 ± 1* [B]<br>5 ± 0* [C] | | |

*Errors calculated by error propagation

**Errors correspond to 68% confidence interval for estimated values from exponential fits

**Supplementary Table SXIII. Results of the Wilcoxon rank-sum tests on the dwell times of a tetrasome (TS) in the left- and right-handed conformation in buffer A.** An h-value of h=0 (h=1) indicates that the data sets – obtained from filtering by averaging over 3.3 s [N=330] – (do not) originate from independent, continuously distributed random samples with equal medians at 5% significance level, i.e. with 95% confidence. The p-value essentially corresponds to the probability of the similarity between the two data sets. Unlike the t-test, this test does not assume normal distributions and is therefore applicable to the exponentially distributed dwell time data (see **Figure 3A-C** in the **main text** and **Figure S8A-C**).

| | | *LEFT-HANDED* | | | RIGHT-HANDED | | |
|---|---|---|---|---|---|---|---|
| | **Buffer A *+/-NAP1*** | **TS on $DNA_{random}$* *+NAP1*** | **TS on $DNA_{w/601}$ +NAP1** | **TS on $DNA_{w/601}$ -NAP1** | **TS on $DNA_{random}$* +NAP1** | **TS on $DNA_{w/601}$ +NAP1** | **TS on $DNA_{w601}$ -NAP1** |
| *LEFT-HANDED* | **TS on $DNA_{random}$* *+NAP1*** | -- | ***h = 1 p = 6.8*10$^{-7}$*** | ***h = 1 p = 7.0*10$^{-3}$*** | h = 1 p = 2.2*10$^{-21}$ | h = 1 p = 1.5*10$^{-14}$ | h = 1 p = 8.7*10$^{-5}$ |
| | **TS on $DNA_{w/601}$ *+NAP1*** | | -- | ***h = 1 p = 3.9*10$^{-14}$*** | h = 1 p = 1.9*10$^{-12}$ | h = 1 p = 1.6*10$^{-5}$ | h = 0 p = 0.47 |
| | **TS on $DNA_{w/601}$ *-NAP1*** | | | -- | h = 1 p = 3.6*10$^{-26}$ | h = 1 p = 2.1*10$^{-22}$ | h = 1 p = 2.2*10$^{-10}$ |
| RIGHT-HANDED | **TS on $DNA_{random}$* *+NAP1*** | | | | -- | **h = 1 p = 5.9*10$^{-4}$** | **h = 1 p = 5.7*10$^{-12}$** |
| | **TS on $DNA_{w/601}$ +NAP1** | | | | | -- | **h = 1 p = 2.1*10$^{-5}$** |
| | **TS on $DNA_{w/601}$ -NAP1** | | | | | | -- |

*data sets obtained by dwell time analysis of the linking number data from our previous study with $(H3.1\text{-}H4)_2$ tetrasomes assembled on DNA molecules with a random sequence (2).

**Supplementary Table SXIV. Results of the Wilcoxon rank-sum tests on the dwell times of a tetrasome in the left- or right-handed state on a DNA molecule with a 601-sequence in different buffers.** An h-value of 0 (1) indicates that the data sets – obtained from filtering by averaging over 3.3 s [N=330] – (do not) originate from independent, continuously distributed random samples with equal medians at 5% significance level, i.e. with 95% confidence. The p-value essentially corresponds to the probability of similarity between the two data sets. Unlike the t-test, this test does not assume normal distributions and is therefore applicable to the exponentially distributed dwell time data (**Figure 3A-C** in **main text**, and **Figure S8A-C**).

RIGHT-HANDED (above the diagonal, right side) / LEFT-HANDED (below the diagonal, left side)

| Buffers + or -*NAP1* | Buffer A +*NAP1* | Buffer B +*NAP1* | Buffer C +*NAP1* | Buffer A -*NAP1* | Buffer B -*NAP1* | Buffer C -*NAP1* |
|---|---|---|---|---|---|---|
| **Buffer A +*NAP1*** | --- | h = 1 p = $5.6*10^{-8}$ | h = 1 p = $8.6*10^{-6}$ | h = 1 p = $2.1*10^{-5}$ | h = 1 p = 0.02 | h = 0 p = 0.37 |
| **Buffer B +*NAP1*** | *h = 1 p = $1.9*10^{-3}$* | --- | h = 0 p = 0.93 | h = 0 p = 0.79 | h = 1 p = $8.4*10^{-3}$ | h = 1 p = $6.1*10^{-16}$ |
| **Buffer C +*NAP1*** | *h = 1 p = $1.6*10^{-37}$* | *h = 1 p = $1.9*10^{-25}$* | --- | h = 0 p = 0.78 | h = 1 p = 0.02 | h = 1 p = $8.1*10^{-10}$ |
| **Buffer A -*NAP1*** | *h = 1 p = $3.9*10^{-14}$* | *h = 1 p = $3.5*10^{-7}$* | *h = 1 p = $1.2*10^{-6}$* | --- | h = 1 p = 0.03 | h = 1 p = $4.2*10^{-9}$ |
| **Buffer B -*NAP1*** | *h = 1 p = $3.7*10^{-46}$* | *h = 1 p = $2.6*10^{-33}$* | *h = 0 p = 0.64* | *h = 1 p = $3.8*10^{-8}$* | --- | h = 1 p = $1.5*10^{-4}$ |
| **Buffer C -*NAP1*** | *h = 1 p = $1.6*10^{-33}$* | *h = 1 p = $6.6*10^{-18}$* | *h = 1 p = $3.0*10^{-9}$* | *h = 0 p = 0.64* | *h = 1 p = $2.5*10^{-12}$* | --- |

**Supplementary Table SXV. Results of the Wilcoxon rank-sum tests on the dwell times of a tetrasome in the left- and right-handed state on a DNA molecule with a 601-sequence in different buffers.** An h-value of 0 (1) indicates that the data sets – obtained from filtering by averaging over 3.3 s [N=330] – (do not) originate from independent, continuously distributed random samples with equal medians at 5% significance level, i.e. with 95% confidence. The p-value essentially corresponds to the probability of similarity between the two data sets. Unlike the t-test, this test does not assume normal distributions and is therefore applicable to the exponentially distributed dwell time data (**Figure 3A-C** in **main text**, and **Figure S8A-C**).

| | | RIGHT-HANDED | | | | | |
|---|---|---|---|---|---|---|---|
| | **Buffers<br>+ or -*NAP1*** | **Buffer A<br>+*NAP1*** | **Buffer B<br>+*NAP1*** | **Buffer C<br>+*NAP1*** | **Buffer A<br>-*NAP1*** | **Buffer B<br>-*NAP1*** | **Buffer C<br>-*NAP1*** |
| LEFT-HANDED | **Buffer A +*NAP1*** | h = 1<br>p =<br>$1.6*10^{-5}$ | h = 0<br>p = 0.51 | h = 0<br>p = 0.63 | h = 0<br>p = 0.47 | h = 0<br>p = 0.07 | h = 1<br>p =<br>$2.8*10^{-10}$ |
| LEFT-HANDED | **Buffer B +*NAP1*** | h = 1<br>p =<br>$7.0*10^{-12}$ | h = 1<br>p =<br>$2.4*10^{-3}$ | h = 1<br>p = 0.02 | h = 0<br>p = 0.05 | h = 1<br>p =<br>$7.0*10^{-6}$ | h = 1<br>p =<br>$9.7*10^{-22}$ |
| LEFT-HANDED | **Buffer C +*NAP1*** | h = 1<br>p =<br>$5.4*10^{-46}$ | h = 1<br>p =<br>$2.8*10^{-42}$ | h = 1<br>p =<br>$8.5*10^{-31}$ | h = 1<br>p =<br>$4.4*10^{-28}$ | h = 1<br>p =<br>$1.1*10^{-39}$ | h = 1<br>p =<br>$1.4*10^{-62}$ |
| LEFT-HANDED | **Buffer A -*NAP1*** | h = 1<br>p =<br>$2.1*10^{-22}$ | h = 1<br>p =<br>$3.2*10^{-15}$ | h = 1<br>p =<br>$1.5*10^{-11}$ | h = 1<br>p =<br>$2.2*10^{-10}$ | h = 1<br>p =<br>$2.9*10^{-17}$ | h = 1<br>p =<br>$2.6*10^{-32}$ |
| LEFT-HANDED | **Buffer B -*NAP1*** | h = 1<br>p =<br>$3.1*10^{-57}$ | h = 1<br>p =<br>$1.4*10^{-53}$ | h = 1<br>p =<br>$2.6*10^{-35}$ | h = 1<br>p =<br>$3.2*10^{-32}$ | h = 1<br>p =<br>$2.5*10^{-47}$ | h = 1<br>p =<br>$4.3*10^{-80}$ |
| LEFT-HANDED | **Buffer C -*NAP1*** | h = 1<br>p =<br>$2.5*10^{-52}$ | h = 1<br>p =<br>$1.9*10^{-40}$ | h = 1<br>p =<br>$2.2*10^{-23}$ | h = 1<br>p =<br>$1.6*10^{-19}$ | h = 1<br>p =<br>$3.2*10^{-37}$ | h = 1<br>p =<br>$2.4*10^{-89}$ |

### SUPPLEMENTARY FIGURES

**A**

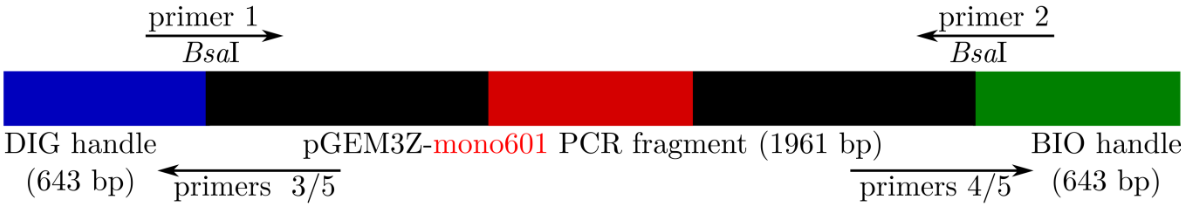

**B**

CCATCTTGGTCTCCtaggtgttgagatccagttcgatgtaacccactcgtgcacccaactgatcttcagcatcttttac
tttcaccagcgtttctgggtgagcaaaaacaggaaggcaaaatgccgcaaaaaagggaataagggcgacacggaaatgttgaa
tactcatactcttcctttttcaatattattgaagcatttatcagggttattgtctcatgagcggatacatatttgaatgtatttagaa
aaataaacaaataggggttccgcgcacatttccccgaaaagtgccacctgacgtctaagaaaccattattatcatgacattaacct
ataaaaataggcgtatcacgaggccctttcgtctcgcgcgtttcggtgatgacggtgaaaacctctgacacatgcagctcccggag
acggtcacagcttgtctgtaagcggatgccgggagcagacaagcccgtcagggcgcgtcagcgggtgttggcgggtgtcggggc
tggcttaactatgcggcatcagagcagattgtactgagagtgcaccatatgcggtgtgaaataccgcacagatgcgtaaggagaa
aataccgcatcaggcgccattcgccattcaggctgcgcaactgttgggaagggcgatcggtgcgggcctcttcgctattacgccag
ctggcgaaagggggatgtgctgcaaggcgattaagttgggtaacgccagggttttcccagtcacgacgttgtaaaacgacggcca
gtgaattgtaatacgactcactatagggcgaattcgagctcggtacccggggatcctctagagtccgggatcctaatgaccaagg
aaagcatgattcttcacaccgagttcatcccttatgtgatggaccctatacgcggccgccctggagaatcccggtgccgaggccgc
tcaattggtcgtagacagctctagcaccgcttaaacgcacgtacgcgctgtcccccgcgttttaaccgccaaggggattactccct
agtctccaggcacgtgtcagatatatacatcctgtgcatgtattgaacagcgaccttgccggtgccagtcggatagtgttccgagc
tcccgacctgcaggcatgcaagcttgagtattctatagtgtcacctaaatagcttggcgtaatcatggtcatagctgtttcctgtgt
gaaattgttatccgctcacaattccacacaacatacgagccggaagcataaagtgtaaagcctggggtgcctaatgagtgagcta
actcacattaattgcgttgcgctcactgcccgctttccagtcgggaaacctgtcgtgccagctgcattaatgaatcggccaacgcgc
ggggagaggcggtttgcgtattgggcgctcttccgcttcctcgctcactgactcgctgcgctcggtcgttcggctgcggcgagcggt
atcagctcactcaaaggcggtaatacggttatccacagaatcaggggataacgcaggaaagaacatgtgagcaaaaggccagc
aaaaggccaggaaccgtaaaaaggccgcgttgctggcgtttttccataggctccgcccccctgacgagcatcacaaaaatcgacg
ctcaagtcagaggtggcgaaacccgacaggactataaagataccaggcgtttccccctggaagctccctcgtgcgctctcctgttc
cgaccctgccgcttaccggatacctgtccgcctttctcccttcgggaagcgtggcgctttctcatagctcacgctgtaggtatctcag
ttcggtgtaggtcgttcgctccaagctgggctgtgtgcacgaaccccccgttcagcccgaccgctgcgccttatccggtaactatcg
tcttgagtccaacccggtaagacacgacttatcgccactggcagcagccactggtaacaggatttgaGGAGACCAAGA
TGG

**Supplementary Figure S1. Specifications of the DNA molecule used in this work. A** Schematics of the 1.96 kbp linear DNA fragments (black) prepared from a pGEM3Z plasmid containing a single 601-sequence (red) in the center (pGEM3Z-mono601) by PCR with primers 1 and 2 (**Table SI**). These main fragments were ligated to a shorter digoxigenin-coated (DIG, blue) fragment (handle) at one end and to a biotin-coated (BIO, green) handle at the other end via *Bsa*I sites. The handles were generated by combinations of primer 5 with primers 3 or 4, respectively (**Table SI**). **B** Sequence of the main DNA fragment containing a single 601-sequence (red) at its center.

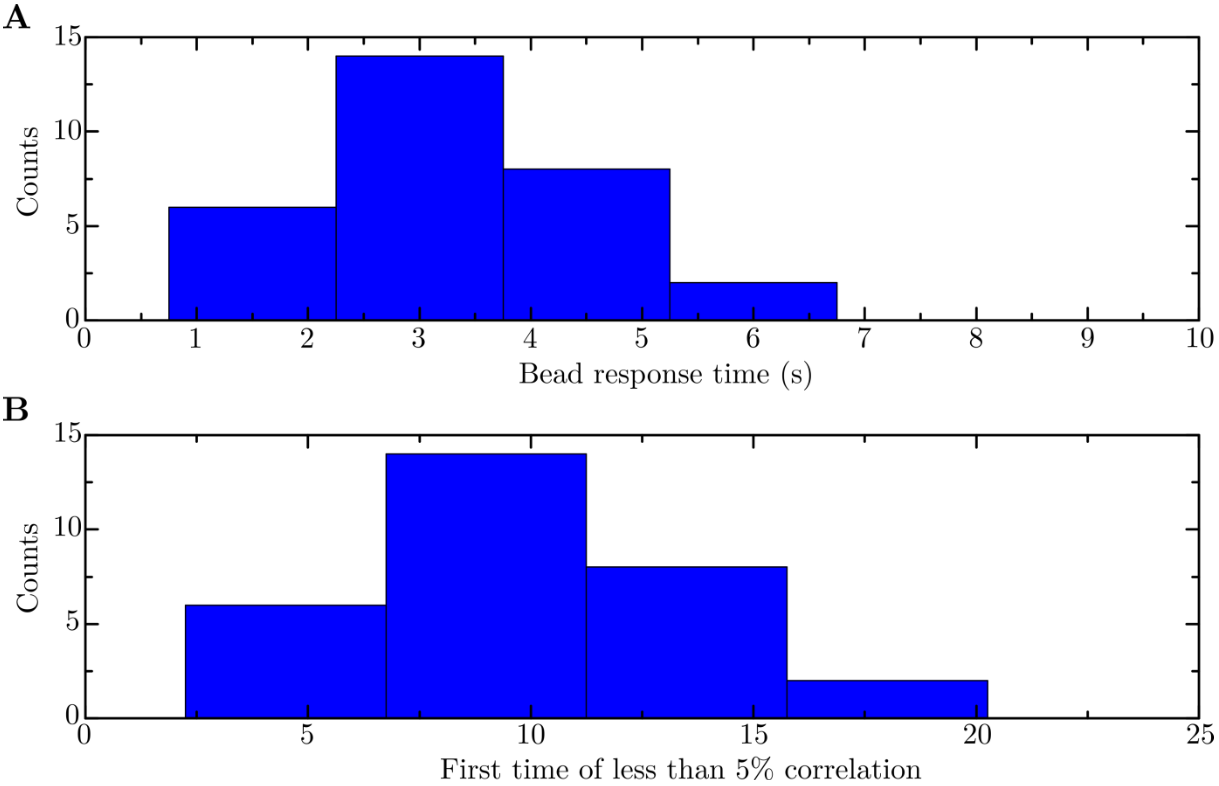

**Supplementary Figure S2. Characteristic times of the bead's angular fluctuations. A** Histogram of the response times of a superparamagnetic bead (**Table SII)** tethered to a 1.96 kbp DNA fragment containing a 601-sequence in its center (**Figure S1)**. The response times are obtained by autocorrelation analysis of the DNA linking number time traces from FOMT measurements as described in Ref. (3). Several measurements ($N$=30) yielded a mean value of $\tau_c$ = 3.3 ± 1.0 s. **B** Histogram of the first times at which the correlation decreases to less than 5%. The data ($N$=30) resulted in a mean value of $\tau_{c,5\%}$ = 9.8 ± 3.1 s. This mean value plus three times its STD ($\tau_{c,long}$ = 19.1 s) was used as an upper boundary for the time difference between steps in the DNA length and DNA linking number time traces to be considered as coinciding.

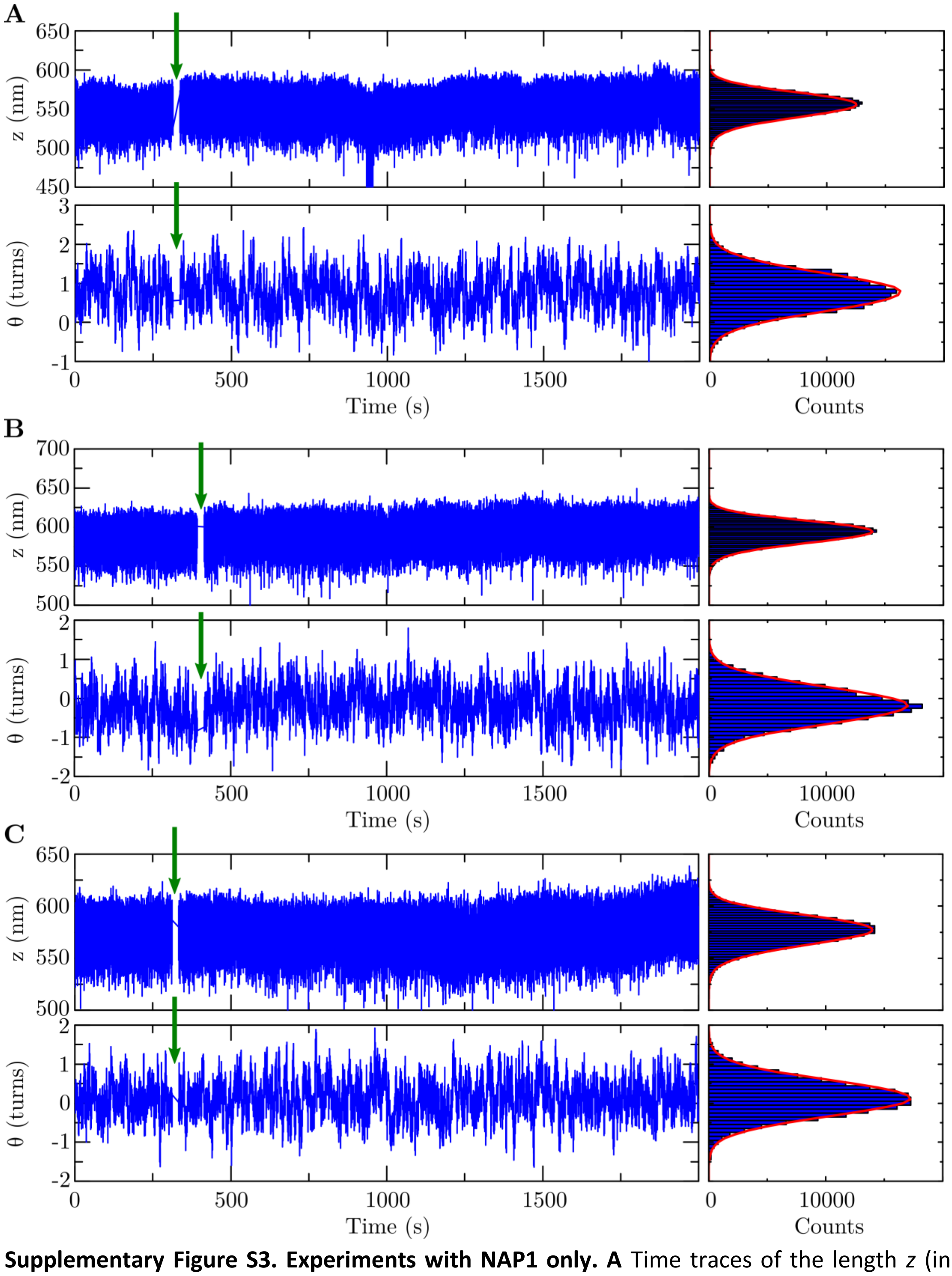

**Supplementary Figure S3. Experiments with NAP1 only. A** Time traces of the length *z* (in nm, top panel) and the linking number *Θ* (in turns, bottom panel) of a DNA molecule before and after flushing in (green arrows) only NAP1 chaperones under the standard conditions of this study in buffer A (see **Materials and Methods** and **Table I** in the **main text**). **B** As in panel (A), but now in buffer B. **C** As in panels (A) and (B), but now in buffer C. As the respective singly peaked mirrored gamma (due to the slight skew of the data to smaller values) and normal distributions (red lines) indicate, NAP1 proteins alone do not interact with the DNA molecule.

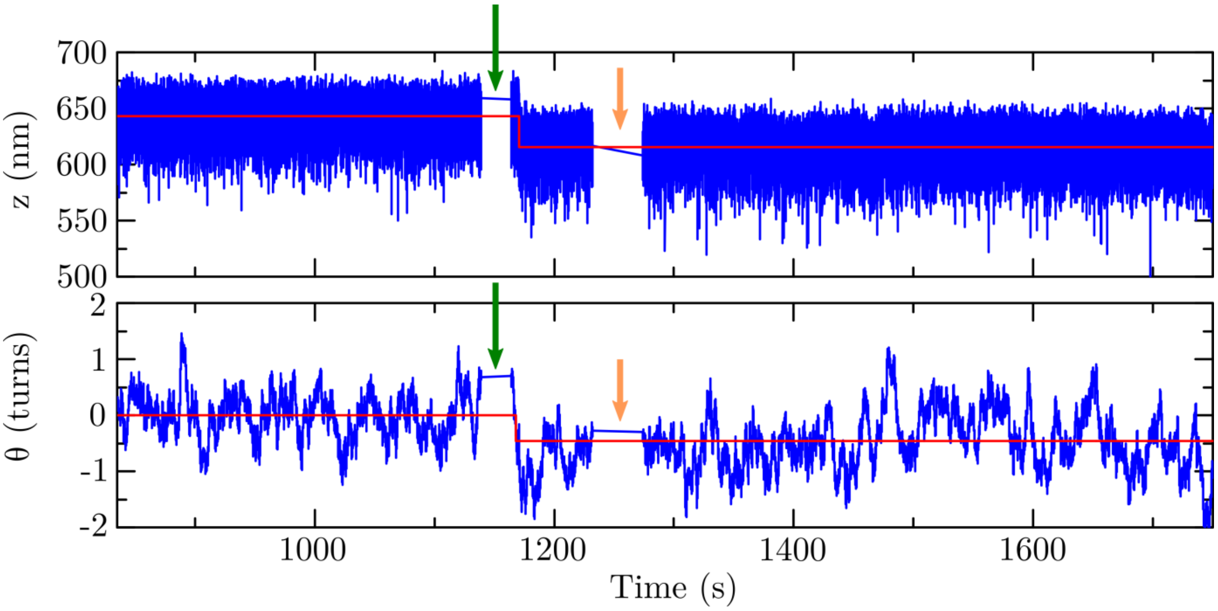

**Supplementary Figure S4. Assembly of a single spontaneously loaded tetrasome onto a DNA molecule with a 601-sequence.** Partial time traces of the length *z* (in nm, top panel) and the linking number *Θ* (in turns, bottom panel) of a $DNA_{w/601}$ before and upon the assembly of a $(H3.1\text{-}H4)_2$ tetrasome without NAP1 in buffer A (see **Table I** in the **main text**). The formation of a tetrasome simultaneously decreased both quantities in the form of a step identified using a step-finder algorithm (red lines) (see **Materials and Methods** in the **main text**). About 60 s after assembly, free proteins were flushed out with measurement buffer (orange arrows) to prevent further histone binding.

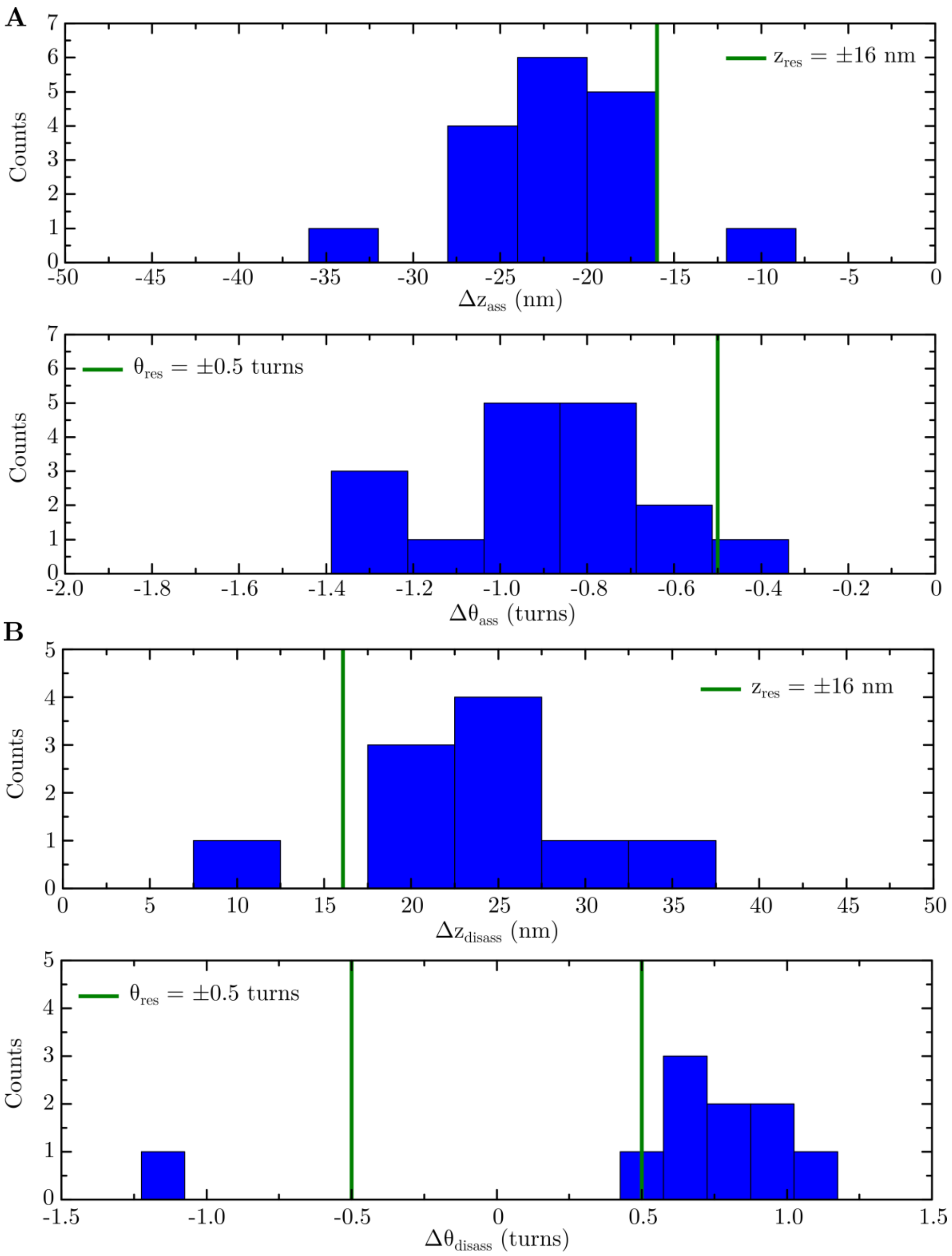

**Supplementary Figure S5. Changes in DNA length and DNA linking number upon assembly and disassembly of single tetrasomes onto DNA molecules with a 601-sequence. A** Histogram of changes in DNA length $\Delta z_{ass}$ (in nm, top panel) and in DNA linking number $\Delta\Theta_{ass}$ (in turns, bottom panel) upon tetrasome assembly in all buffer conditions (see **Table SIII**). **B** Histogram of changes in DNA length $\Delta z_{disass}$ (in nm, top panel) and in DNA linking number $\Delta\Theta_{disass}$ (in turns, bottom panel) upon tetrasome disassembly in all buffer conditions (see **Table SIV**). These datasets include data from both NAP1-loaded and spontaneously loaded tetrasomes (see main text)**.** The green lines depict the mean spatial resolutions based on 1 average standard deviation (1 STD = ±16 nm and 1 STD = ±0.5 turns) determined from all experiments (see **Materials and Methods** in the main text). The data within resolution yielded a mean value of $\Delta z_{ass}$ = -22 ± 5 nm/$\Delta\Theta_{ass}$ = -0.9 ± 0.2 turns and $\Delta z_{disass}$ = 25 ± 5 nm/ $\Delta\Theta_{disass}$ = 0.8 ± 0.2 turns (considering the absolute value of the single negative value indicating right-handed disassembly), respectively.

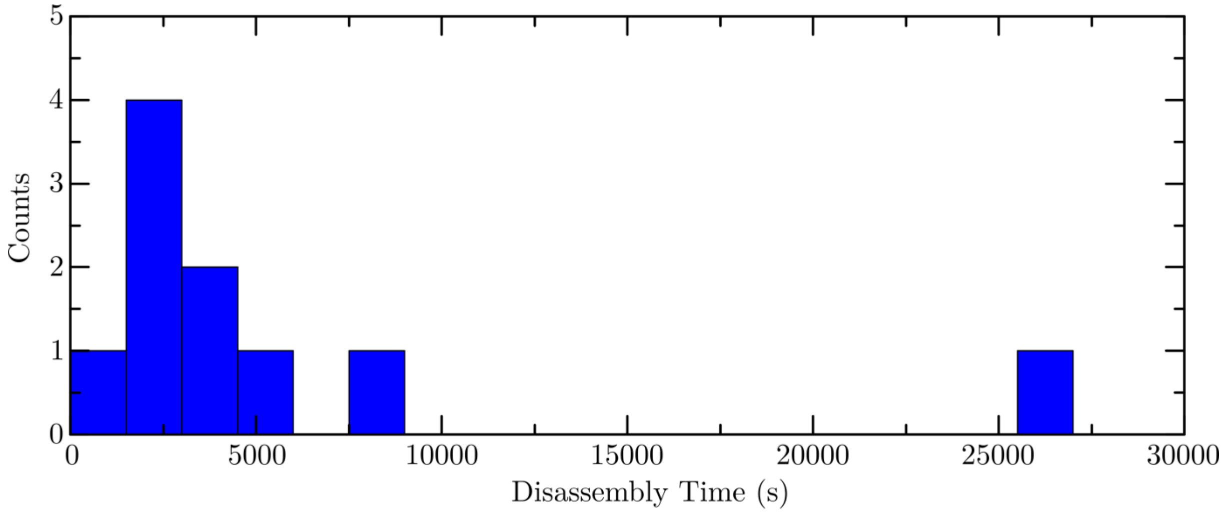

**Supplementary Figure S6. Distribution of the disassembly times for single tetrasomes from DNA molecules with a 601-sequence.** This dataset includes data from both NAP1-loaded and spontaneously loaded tetrasomes (see main text)**.** In 40% (*n*=8) of all experiments (*N*=20), the single tetrasomes assembled on DNA molecules with a centered 601-sequence were observed to disassemble in the course of the measurements at different times (**Table SIV** and **Figure S5**). The data yielded a mean value of $\tau_{disass}$ = 3364 ± 765 s (1 standard error of the mean (SEM) due to broadness). The longest disassembly time at 26487 s was excluded from this calculation, as it is even longer than the mean duration $t_m$ = 22665 ± 3141 s (1 SEM) of the measurements. The dissociation times can be critical, but are much longer than the time scales of the relevant dynamics, such as the dwell times, investigated in this study (see **Materials and Methods**, and **Figure 3A-C** in the **main text,** and **Figure S8A-C)**.

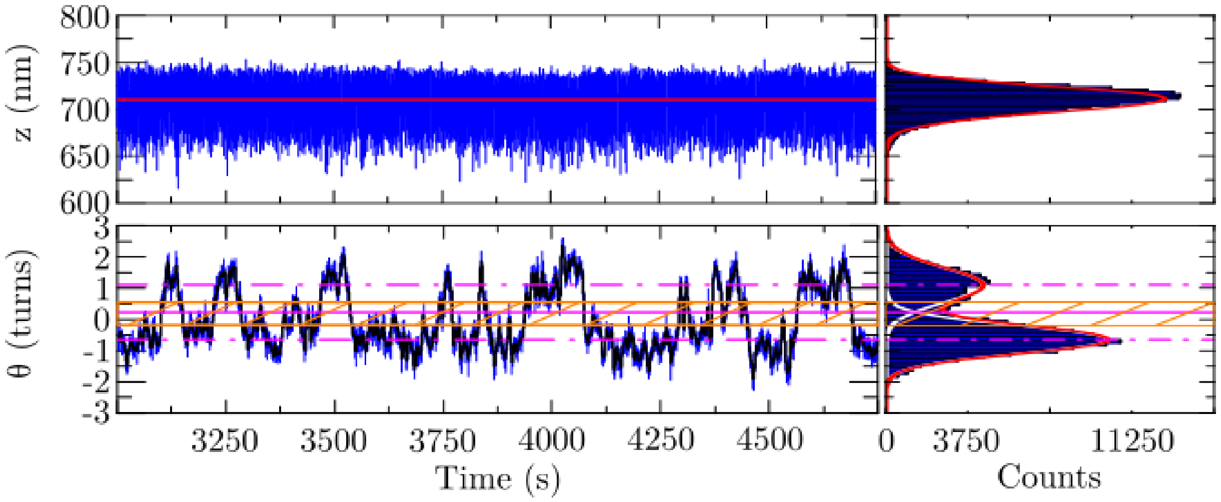

**Supplementary Figure S7. Flipping of a single spontaneously loaded tetrasome on a DNA molecule with a 601-sequence.** Partial time traces of the length *z* (in nm, top panel) and the linking number *Θ* (in turns, bottom panel) of a DNA molecule after the assembly of a $(H3.1\text{-}H4)_2$ tetrasome in buffer A (see **Table I**) upon flushing in histone tetramers only. As indicated from the fit by the step-finder algorithm to the time trace (red line, left panel) and the fit of a mirrored gamma function (red line in histogram plot, right panel) to the skewed data, the DNA length stays constant. The DNA linking number spontaneously fluctuates, i.e. 'flips', between two states identified by fitting two Gaussian functions (white lines in histogram plot, right panel) underlying the full profile (red line in histogram plot, right panel). The two states correspond to a prevalent left-handed and a less adopted right-handed conformation of DNA wrapping with the respective mean linking numbers $\Theta_{left}$ = -0.65 ± 0.02 turns and $\Theta_{right}$ = +1.10 ± 0.04 turns (dashed-dotted magenta lines, 95% confidence level for estimated values). These structural dynamics were quantified in terms of the dwell times in the two states based on a threshold zone (shaded orange area) set by 1 STD from each mean value (orange solid lines) around their average (solid magenta line) (see **Materials and Methods** in the **main text**).

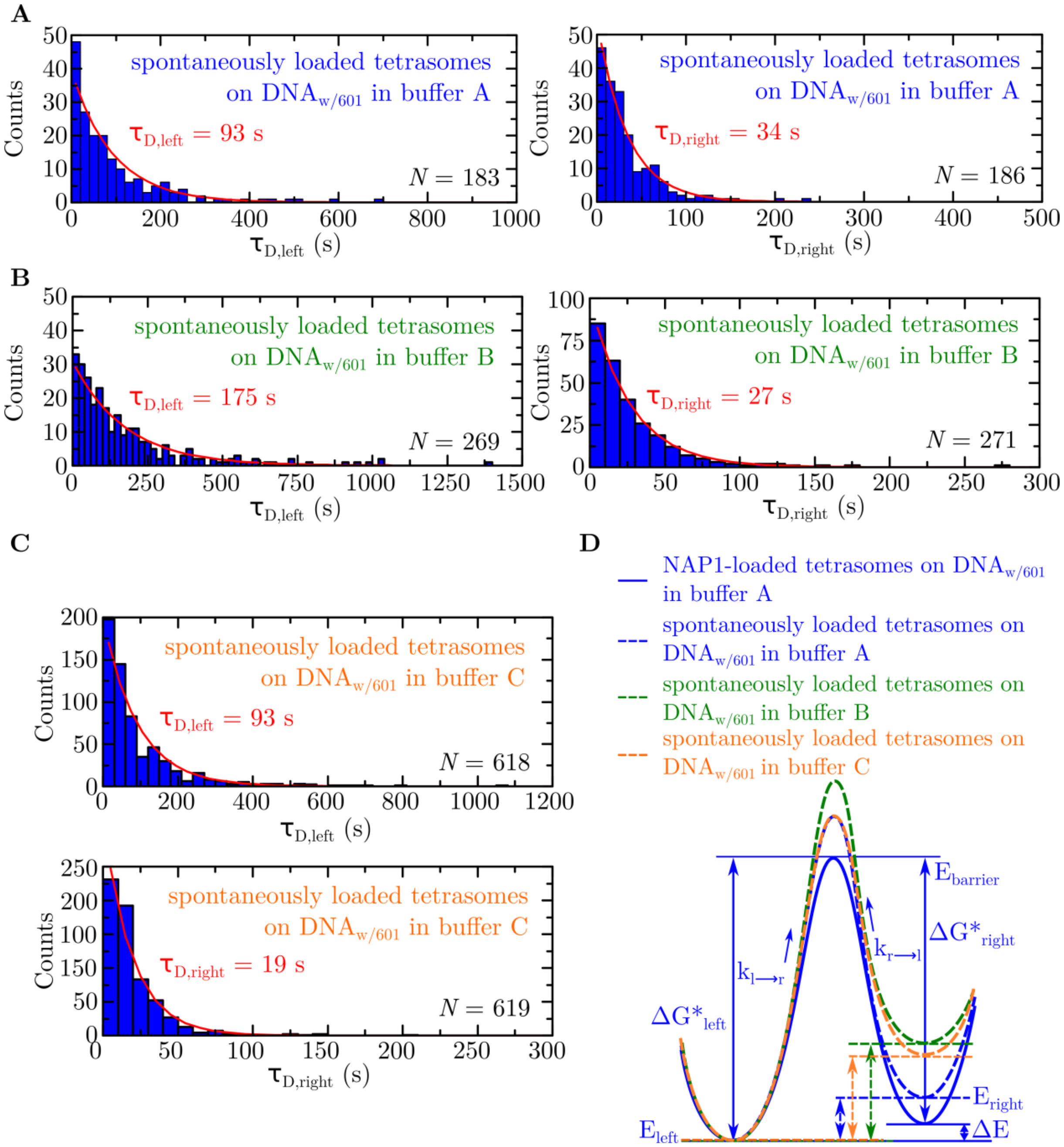

**Supplementary Figure S8. Dwell times of single spontaneously loaded tetrasomes on DNA molecules with a 601-sequence. A** Histograms of the dwell times of a single $(H3.1\text{-}H4)_2$ tetrasome loaded on a $DNA_{w/601}$ molecule in the absence of NAP1 ('spontaneously loaded tetrasomes') in the left- (left panel) and right-handed conformation (right panel) in buffer A (**Table I** in the **main text** and **Table SXII**). Exponential fits (red lines) yield mean dwell times $\tau_{D,left}$ = 93 +7/-6 s (*N*=183) and $\tau_{D,right}$ = 34 +3/-2 s (*N*=186), respectively (68% confidence level for estimated values). **B** As in (A), but now in buffer B (**Table I** in the **main text** and **Table SXII**). Exponential fits (red lines) yielded a mean dwell time of $\tau_{D,left}$ = 175 +11/-10 s (*N*=269) and $\tau_{D,right}$ = 27 ± 2 s (*N*=271), respectively (68% confidence level for estimated values). **C** Histograms of the dwell times of a single self-loaded $(H3.1\text{-}H4)_2$ tetrasome on a $DNA_{w/601}$ molecule in the left- (top panel) and right-handed conformation (bottom panel) in buffer C (**Table I** in the **main text** and **Table SXII**). Exponential fits (red lines) yielded a mean dwell time of $\tau_{D,left}$ = 93 ± 4 s (*N*=618) and $\tau_{D,right}$ = 19 ± 1 s (*N*=619), respectively (68% confidence level for estimated values). All data in panels (A)-(C) were obtained by dwell-time analysis of the corresponding DNA linking number time traces filtered by averaging over 3.3 s (*N*=330) (**Materials and Methods** in the **main text**). **D** Schematic energy diagrams of single spontaneously loaded (H3.1-H4)$_2$ tetrasomes on DNA$_{w/601}$ in all buffer conditions, based on the dwell-time values and the probabilities obtained from the linking number distributions (**Tables SXI-SXV**). The free energy differences ($\Delta E$) between the left- and right-handed conformations of spontaneously loaded tetrasomes in buffer A (dashed blue lines), with the respective energies $E_{left}$ and $E_{right}$, are considerably increased compared to NAP1-loaded tetrasomes in the same buffer A (**Table II** and **Figure 3** in the **main text**). For spontaneously loaded tetrasomes in buffer B (dashed green lines), $\Delta E$ is largest, following that for spontaneously loaded tetrasomes in buffer A, as well as in buffer C (dashed orange lines). The heights of the energy barriers $\Delta G^*_{left}$ and $\Delta G^*_{right}$ are estimated from the rates $k_{l\text{->}r}$ and $k_{r\text{->}l}$. $\Delta G^*_{left}$ is very similar for spontaneously loaded tetrasomes in buffers A and C, but considerably higher for tetrasomes in buffer B, while $\Delta G^*_{right}$ is essentially unchanged.
